# Megalencephalic leukoencephalopathy with subcortical cysts is a developmental disorder of the gliovascular unit

**DOI:** 10.1101/2021.05.17.444434

**Authors:** Alice Gilbert, Xabier Elorza-Vidal, Armelle Rancillac, Audrey Chagnot, Mervé Yetim, Vincent Hingot, Thomas Deffieux, Anne-Cécile Boulay, Rodrigo Alvear-Perez, Salvatore Cisternino, Sabrina Martin, Sonia Taib, Antoinette Gelot, Virginie Mignon, Maryline Favier, Isabelle Brunet, Xavier Declèves, Mickael Tanter, Raul Estevez, Denis Vivien, Bruno Saubaméa, Martine Cohen-Salmon

**Author notes:** Corresponding author: Martine Cohen-Salmon, Collège de France, Center for Interdisciplinary Research in Biology (CIRB), CNRS, UMR 7241, INSERM, U1050, 11 place Marcelin Berthelot, 75005 Paris, France. coauthors (Armelle Rancillac, Audrey Chagnot, Mervé Yetim).

## Abstract

Absence of the astrocyte-specific membrane protein MLC1 is responsible for megalencephalic leukoencephalopathy with subcortical cysts (MLC); this rare type of leukodystrophy is characterized by early-onset macrocephaly and progressive white matter vacuolation that lead to ataxia, spasticity, and cognitive decline. During postnatal development (from P5 to P15 in the mouse), MLC1 forms a membrane complex with GlialCAM (another astrocytic transmembrane protein) at the junctions between perivascular astrocytic processes (PvAPs, which along with blood vessels form the gliovascular unit (GVU)). We analyzed the GVU in the *Mlc1* knock-out mouse model of MLC. The absence of MLC1 led to an accumulation of fluid in the brain but did not modify the endothelial organization or the integrity of the blood-brain barrier. From P10 onward, the postnatal acquisition of vascular smooth muscle cell contractility was altered, resulting in a marked reduction in arterial perfusion and neurovascular coupling. These anomalies were correlated with alterations in astrocyte morphology, astrocyte polarity and the structural organization of the PvAP’s perivascular coverage, and poor intraparenchymal circulation of the cerebrospinal fluid (CSF). Hence, MLC1 is required for the postnatal development and organization of PvAPs and controls vessel contractility and intraparenchymal interstitial fluid clearance. Our data suggest that (i) MLC is a developmental disorder of the GVU, and (ii) PvAP and VSMC maturation defects are primary events in the pathogenesis of MLC and therapeutic targets for this disease.

## Introduction

Megalencephalic leukoencephalopathy with subcortical cysts (MLC) is a rare type of leukodystrophy (OMIM 604004) mainly caused by mutations in the *MLC1* gene (MIM #605908) [1, 2]. Patients with MLC display early-onset macrocephaly and progressive white matter vacuolation, leading to slowly progressive ataxia, spasticity, and cognitive decline. Most mutations in *MLC1* result in the degradation of the encoded protein MLC1 [3, 2, 4], a membrane protein that is specifically expressed by the astrocytic lineage in the brain and present at high levels at the junctions between perivascular astrocytic processes (PvAPs) [5, 6]. At present, there is no cure for MLC - only symptomatic treatments and supportive care are available. Although the physiopathological mechanisms leading to MLC have not been characterized, the strong expression of MLC1 in PvAPs and other recent observations suggest that the protein has a role in gliovascular functions, in particular the regulation of ion/water homeostasis [7, 8, 5]. Indeed, MLC patients present widespread brain edema and swollen PvAPs [9, 10]. GlialCAM, another transmembrane protein forming a complex with MLC1 and responsible for its endoplasmic reticulum exit, is an auxiliary subunit of ClC-2, an inward rectifier chloride channel expressed in a subtype of astrocytes [11, 5, 10, 12, 13]. Recessive and dominant mutations in GlialCAM cause MLC subtypes MLC2A and MLC2B respectively [14]. *In vitro*, the MLC1/GlialCAM complex indirectly regulates other ion channels, such as TRPV4 and LRRC8 [15].

Despite the above observations, it is still not known whether and how MLC1 influences the physiology of the gliovascular unit (GVU, the functional interface comprising PvAPs and the brain vessels). We recently reported that MLC1 expression in mouse PvAPs starts around postnatal day (P)5 and that the MLC1/GlialCAM complexes in PvAPs form progressively from P5 to P15, with the protein deposits creating a meshwork between the astrocytes’ perivascular membranes [16]. We also demonstrated that this postnatal period is a developmental window for the molecular maturation of brain endothelial cells (ECs), particularly with regard to their efflux properties and for the contractility of vascular smooth muscle cells (VSMCs) [16, 17]. Given that astrocytes are key regulators of cerebrovascular development and function (e.g. BBB integrity, immune quiescence and perivascular homeostasis, neurovascular coupling) [18–21], we hypothesized that the MLC1/GlialCAM complex might influence the postnatal differentiation of the vascular system.

Here, we characterized key aspects of the molecular and functional organization of the GVU in *Mlc1* KO mice, a preclinical model of MLC that recapitulates several important features of the disease and that can be used to examine the pathological cascade [5]. Our results revealed that MLC1 is a critical factor in the postnatal maturation and function of the GVU.

## Materials and Methods

### Animals

All animal experiments were carried out in compliance with the European Directive 2010/63/EU on the protection of animals used for scientific purposes and the guidelines issued by the French National Animal Care and Use Committee (reference: 2019021814073504 and 2019022113258393). *Mlc1* KO mice were maintained on a C57BL6 genetic background.

### MVs purification

MVs were isolated from whole brain using selection filtration, as described previously [22]. We purified vessels that passed through 100 µm pores but not 20 µm pores [22]. Brain vessels from two animals were pooled for the 2-month sample, with 3 for P15 and 5 for P5.

### Immunohistochemical analysis of brain slices

Mice were anesthetized with pentobarbital (600 mg/kg, i.p.) and killed by transcardiac perfusion with PBS/PFA 4%. The brain was removed and cut into 40-µm-thick sections using a Leitz (1400) cryomicrotome. Slices were immersed in the blocking solution (PBS/normal goat serum (NGS) 5%/Triton X-100 0.5%) for 1 h at room temperature (RT) and then incubated with primary antibodies (**Table S11**) diluted in the blocking solution 12 h at 4 °C. After 3 washes in PBS, slices were incubated for 2 h at RT with secondary antibodies and Hoechst dye, rinsed in PBS, and mounted in Fluormount G (Southern Biotech, Birmingham, AL).

### Western blot

Proteins were extracted from one brain hemisphere or from purified MVs in 2% SDS (500 µl or 50µl per sample, respectively) with EDTA-free Complete Protease Inhibitor (Roche), sonicated three times at 20 Hz (Vibra cell VCX130) and centrifuged for 20 min at 10,000 g at 4 °C. Supernatants were heated in Laemmli loading buffer for 5 min at 56 °C. Proteins were extracted from one brain hemisphere per sample in 500 µL SDS 2%, under the same conditions. The protein content was measured using the Pierce 660 nm protein assay (Thermo Scientific, Waltham, MA, USA). Equal amounts of proteins were separated by denaturing electrophoresis on Mini-Protean TGX stain-free gels (Biorad) and then electrotransferred to nitrocellulose membranes using the Trans-blot Turbo Transfer System (Biorad). Membranes were hybridized, as described previously [23]. The antibodies used in this study are listed in **Table S11**. Horseradish peroxidase activity was visualized using enhanced chemiluminescence in a Western Lightning Plus system (Perkin Elmer, Waltham, MA, USA). Chemiluminescent imaging was performed on a FUSION FX system (Vilber, South Korea). At least four independent samples were analyzed in each experiment. The level of chemiluminescence for each antibody was normalized against that of a stain-free membrane, or histone H3.

### Quantitative RT-PCR

RNA was extracted using the Rneasy Lipid Tissue Mini Kit (Qiagen, Hilden, Germany). cDNA was then generated using the Superscript™ III Reverse Transcriptase Kit (Thermo Fisher). Differential levels of cDNA expression were measured using droplet digital PCR. Briefly, cDNA and primers (**Table S11**) were distributed into approximately 10,000 to 20,000 droplets. cDNAs were then PCR-amplified in a thermal cycler and read (as the number of positive and negative droplets) with a QX200 Droplet Digital PCR System (Biorad). The ratio for each tested gene was normalized against the total number of positive droplets for *Gapdh*.

### *In situ* brain perfusion

Mice were anesthetized with ketamine-xylazine (140 and 8 mg/kg, respectively, i.p.), and a polyethylene catheter was inserted into the carotid veins. The heart was incised, and the perfusion was started immediately (flow rate: 2.5 mL/min) so as to completely replace the blood with Krebs carbonate-buffered physiological saline (128 mM NaCl, 24 mM NaHCO_3_, 4.2 mM KCl, 2.4 mM NaH_2_PO_4_, 1.5 mM CaCl_2_, 0.9 mM MgCl_2)_, 9 mM D-glucose) supplemented with [^14^C] sucrose (0.3 µCi/mL) (Perkin Elmer Life Sciences, Courtaboeuf, France) as a marker of vascular integrity. The saline was bubbled with 95% O_2_/5% CO_2_ for pH control (7.4) and warmed to 37 °C. Perfusion was terminated after 120 sec by decapitating the mouse. The whole brain was removed from the skull and dissected out on a freezer pack. Brain hemisphere and two aliquots of perfusion fluid were placed in tared vials and weighed, digested with Solvable^®^ (Perkin Elmer) and mixed with Ultima gold XR^®^ (Perkin Elmer) for ^14^C dpm counting (Tri-Carb^®^, Perkin Elmer). In some experiments, human serum albumin (40 g/L) (Vialebex®, Paris, France) was added to the perfusion fluid in order to increase the hydrostatic pressure (∼180 mmHg) and create shear stress [23]. The brain [^14^C]-sucrose vascular volume (Vv, in µL/g) was calculated from the distribution of the [^14^C]-sucrose: Vv = X_v_/C_v_ where X_v_ (dpm/g) is the [^14^C] sucrose concentration in the hemispheres and C_v_ (dpm/µL) is the [^14^C] sucrose concentration in the perfusion fluid [24]. It should be noted that in mammals, the very hydrophilic, low-molecular-weight (342 Da) disaccharide sucrose does not bind to plasma proteins and does not have a dedicated transporter. Accordingly, sucrose does not diffuse passively and thus serves as a marker of BBB integrity [25]. In this context, variations in sucrose’s distribution volume in the brain solely reflect changes in the BBB’s physical integrity.

### VSMC responsiveness

Mice were rapidly decapitated, and the brains were quickly removed and placed in cold (∼4 °C) artificial cerebrospinal fluid (aCSF) solution containing 119 mM NaCl, 2.5 mM KCl, 2.5 mM CaCl2, 26.2 mM NaHCO3, 1 mM NaH2PO4, 1.3 mM MgSO4, 11 mM D-glucose (pH = 7.35). Brains were constantly oxygenated with 95% O_2_ –5% CO_2_. Brain cortex slices (400 µm thick) were cut with a vibratome (VT2000S, Leica) and transferred to a constantly oxygenated (95% O_2_–5% CO_2_) holding chamber containing aCSF. Subsequently, individual slices were placed in a submerged recording chamber maintained at RT under an upright microscope (Zeiss) equipped with a CCD camera (Qimaging) and perfused at 2 ml/min with oxygenated aCSF. Only one vessel per slice was selected for measurements of vascular responsiveness, at the junction between layers I and II of the somatosensory cortex and with a well-defined luminal diameter (10–15 µm). An image was acquired every 30 s. Each recording started with the establishment of a control baseline for 5 min. Vessels with an unstable baseline (i.e. a change in diameter of more than 5%) were discarded from analysis. Vasoconstriction was induced by the application of the thromboxane A_2_ receptor agonist U46619 (9,11-dideoxy-11a,9a-epoxymethanoprostaglandin F2α, 50 nM, Sigma) for 2 min. The signal was recorded until it had returned to the baseline.

### Functional ultrasound

Two-month-old mice were anesthetized, and CBF responses to whisker stimulation were determined using fUS. The protocol is described in detail in [26]. Briefly, mice were intubated and mechanically ventilated (frequency: 120/min; Tidal volume:10 ml/kg) by maintaining anesthesia with 2% isoflurane in 70% N_2_O/30% O_2_. Mice were placed in a stereotaxic frame, and the head was shaved and cleaned with povidone-iodine. An incision was made along the midline head skin (to expose the skull), and lidocaine spray was applied to the head. Whiskers on the left side were cut to a length of one centimeter. Anesthesia was switched to a subcutaneous infusion of medetomidine (Domitor®, Pfizer, 0.1 mg/kg) and isoflurane, N_2_O and O_2_ were withdrawn 10 min later. So that the isoflurane could dissipate and the CBF could stabilization, the fUS measurements were initiated 20 min later. Ultrasound gel was applied between the ultrasound probe and the mouse’s skull, to ensure good acoustic coupling.

The probe was positioned in the coronal plane, corresponding to the somatosensory barrel field cortex (S1bf; bregma −1.5 mm). Ultrafast acquisition was performed with an ultrasound sequence based on compounded plane wave transmission (11 angles from 10° to 10°, in increments of 2°), using a 15 MHz probe (Vermon, France; 100 µm x 100 µm in-plane pixels; slice thickness: 300 µm; elevation focus: 8 µm; frame rate: 500 Hz). The whiskers were mechanically stimulated three times for 30 s, interspaced with a 60 s rest period (total duration of the experiment: 300s). Using MATLAB (the MathWorks Inc., Natick, Massachussets, United States), we calculated the coefficient for the correlation between the normalized power Doppler (PD) intensity over time and a step function following the stimulation pattern. An activation map was reconstructed by selecting only pixels with a correlation coefficient above 0.2. The relative PD increase was quantified as the mean PD signal in the activated area.

### Ultrasound localization microscopy

The acquisition and post processing steps for ultrasound localization microscopy were adapted from [27]. For each image, 100 µL of Sonovue microbubbles were injected into the tail vein. Blocks of 800 compounded frames (−5° 0° 5°) at 1 kHz were acquired for 800 ms and saved for 200 ms; this scheme was repeated for 180 s. A combination of a Butterworth high-pass filter (second order, 20 Hz) and a singular value decomposition filter (10 values) was used to separate microbubble echoes from tissue echoes. The microbubbles’ centroid positions were localized using a weighted average algorithm. Microbubbles were tracked through consecutive frames using MathWorks (the MathWorks Inc., Natick, Massachussets, United States). The tracks were interpolated and smoothed using a 5-point sliding window, and redundant positions were removed. A density image was reconstructed on an 11 µm × 10 µm grid.

### Magnetic resonance imaging

MRI was performed on a 7T Pharmascan MRI system (Bruker, Rheinstetten, Germany) equipped with volume transmit and surface receive coils and operated via Paravision^®^ 6.0 software (Bruker, Rheinstetten, Germany). An anatomical T2-weighted acquisition was performed prior to contrast injection, with the following parameters: echo time (TE) = 40 ms; repetition time (TR) = 3500 ms; flip angle (FA) = 90°; averages = 2; number of echoes = 8; on a 256×256 sagittal matrix with 20 contiguous 0.5 mm thick slices (in-plan resolution = 0.07 x 0.07 mm) for a total duration of 2 min 41 s. The ADC was calculated from a multi-b diffusion-weighted echo planar imaging sequence, with following parameters: TE = 35 ms; TR = 2000 ms; FA = 90°; number of segments = 4; 16 b-values = 20 s.mm^-2^, 30 s.mm^-2^, 40 s.mm^-2^, 50 s.mm^-2^, 75 s.mm^-2^, 100 s.mm^-2^, 150 s.mm^-2^, 200 s.mm^-2^, 300 s.mm^-2^, 400 s.mm^-2^, 500 s.mm^-2^, 750 s.mm^-2^, 1000 s.mm^-2^, 1250 s.mm^-2^, 1500 s.mm^-2^, 2000 s.mm^-2^; 12 directions; on a 128×40 sagittal matrix with 9 contiguous 1 mm thick slices (in-plan resolution: 0.15 x 0.15 mm), with use of a saturation band to remove the out-of-matrix signal, over a total duration of 25 min 45 s. The ADC acquisition was performed prior to contrast injection. To estimate the concentration of contrast agent, T1 maps before and after contrast injection were computed from a FAIR RARE (flow alternating inversion recovery, rapid acquisition with refocused echoes) acquisition derived from the Look-Locker T1 mapping sequence [28], with the following parameters: TE = 5.3 ms; TR = 5000 ms; FA = 90°; RARE factor = 4; 10 inversion times (TI) = 10 ms, 21 ms, 44 ms, 195 ms, 410 ms, 862 ms, 1811 ms, 3807 ms, 8000 ms; on a 64×64 single mediosagittal slice (in-plan resolution: 0.3 x 0.3 mm; thickness: 0.8 mm) for a total duration of 4 min 25 s. A single acquisition was performed before contrast injection (to map the reference T1), and 8 consecutive acquisitions were performed 10 minutes after contrast injection – providing dynamic data over 40 minutes.

### Injection of contrast agent into the cerebrospinal fluid

1 µl of 500 mM DOTA-Gd (Dotarem^®^, Guerbet^®^, France) was injected over 1 minute into the CSF with a glass micropipette through the cisterna magna, as described previously [29]. Briefly, the mice were anesthetized with isoflurane (induction: 5%, maintenance: 2-3%) in 70% N_2_O/30% O_2_. The neck was shaved, and lidocaine was sprayed on for local analgesia. A vertical incision was performed, and the muscle planes were separated vertically upon reaching the cisterna magna. A micropipette formed from an elongated capillary glass tube and filled with 1 µL of DOTA-Gd was inserted into the cisterna magna. Before and after injection, one-minute pauses enabled the CSF pressure to normalize. Before micropipette removal, a drop of superglue was added to form a seal and prevent subsequent leakage of CSF. The incision was cleaned and then closed with 5.0 gauge surgical silk thread.

### MRI analyses

T1 values, quantitative contrast measurements and ADC maps were calculated with in-house MATLAB code (R2021a, Natick, Massachusetts: the MathWorks Inc; 2020). Regions of interest were determined using the FIJI image analysis suite [30].

T1 maps were computed from the FAIR RARE data. Briefly, T1 was extracted after fitting the signal recovery equation:

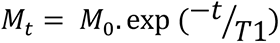

The contrast agent concentration [CA] was determined from the equation:

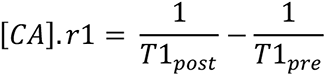

where r1, T1_post_ and T1_pre_ were respectively the T1 relaxivity, the T1 value after contrast, and the T1 value before contrast. The ADC was calculated as the slope of the log of signal loss for the b-value, according to the following equation:

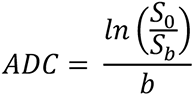

where b = 1000 s.mm^-2^.

### Tissue clearing and immunohistochemical staining

Mice were killed with pentobarbital (600 mg/kg, i.p.). Brains were removed and post-fixed in 4% paraformaldehyde (PFA) for 24 h at 4 °C and then assessed using the “immunolabeling-enabled three-dimensional imaging of solvent-cleared organs” technique [31]. The samples were first dehydrated with increasingly concentrated aqueous methanol solutions (MetOH: 20%, 40%, 60%, 80%, and twice 100%, for 1 h each) at RT and then incubated in 66% dichloromethane (DCM, Sigma Aldrich)/33% MetOH overnight. After 2 washes in 100% MetOH, brains were incubated in 5% H_2_O_2_/MetOH overnight at RT, rehydrated with increasingly dilute aqueous methanol solutions (80%, 60%, 40%, and 20%; 1h each). Before immunostaining, brains were permeabilized first for 2 x 1h at RT in 0.2% Triton X-100/PBS, for 24 h at 37 °C in 0.16% Triton X-100/2.3% glycine/20% DMSO/PBS, and then for 2 days at 37 °C in 0.16% Triton X-100/6% donkey serum/10% DMSO/PBS. Brains were incubated for 3 days at 37 °C with primary antibody diluted in a 0.2 Tween/1% heparin/3% donkey serum/5% DMSO/PBS solution, washed 5 times during 24h at 37 °C in 0.2% Tween20/1% heparin/PBS solution, incubated for 3 days at 37 °C with secondary antibody diluted in a 0.2 Tween/1% heparin/3% donkey serum/PBS solution, and another washed five times. The brain samples were then dehydrated again with a MetOH/H_2_O series (20%, 40%, 60%, 80% and 100% for 1h each, and then 100% overnight) at RT. On the following day, brains were incubated for 3h in 66% DCM/33% MetOH and then twice for 15 min at RT in 100% DCM and lastly cleared overnight in dibenzyl ether.

The cleared tissues were imaged using a light sheet microscope and Inspector pro software (Lavision Biotec GmbH, Bielefeld, Germany). 3D reconstructions of the somatosensory cortex (a 400 µm-thick column for Pecam-1 and 500 to 750 µm for SMA) were visualized with Imaris software (Bitplane). The length and number of branch points of Pecam-1- or SMA-immunolabeled brain vessels were quantified using the “Surface” and “Filament” tools in Imaris software (Oxford instruments, Oxford). Anastomoses were measured by eye.

### Astrocyte morphology

Hippocampal slices were pictured using a 40X objective on a Zeiss Axio-observer Z1 with a motorized XYZ stage (Zeiss, Oberkochen, Germany). To analyze astrocyte ramifications, we adapted a previously described technique [32]. Using ImageJ software, 7 concentric circles at 5 μm intervals were drawn around each astrocyte on confocal Z-stack images. The number of intersections of GFAP-positive astrocytic processes with each circle was counted. We analyzed the astrocytes’ orientation by adapting a previously described technique [33]. Using ImageJ software, a grid delimitating 100 µm^2^ squares was drawn on confocal Z-stack images oriented with the PL or a vessel. The number of intersections of astrocytic GFAP positive processes with horizontal lines (i.e. processes perpendicular to the pyramidal layer or vessel, so-called axial processes) and vertical lines (i.e. processes parallel to the pyramidal layer or vessel, so-called lateral processes) were counted. The cell’s polarity index was defined as the ratio between the axial processes and the lateral processes. A polarity index of 1 indicates no polarity, whereas a polarity index greater than 1 indicates preferentially perpendicular orientation toward the pyramidal layer or the vessel.

### Electron microscopy

Mice were anesthetized with ketamine-xylazine (140 and 8 mg/kg, respectively, i.p.) and transcardially perfused with the fixative (2% paraformaldehyde, 3% glutaraldehyde, 3mM CaCl_2_ in 0.1M cacodylate buffer pH 7.4) for 12 min. The brains were removed and left overnight at 4 °C in the same fixative. Brain fragments (0.3 x 1 x 1 mm^3^) were postfixed first in 0.1M cacodylate buffer pH 7.4 + 1% OsO_4_ for 1h at 4 °C and then in 1% aqueous uranyl acetate for 2h at RT. After dehydration in graded ethanol and then propylene oxide, the fragments were embedded in EPON resin (Electron Microscopy Sciences, Hatfield, PA). Ultrathin (80 nm) sections were prepared, stained with lead citrate and imaged in a Jeol 100S transmission electron microscope (Jeol, Croissy-sur-Seine, France) equipped with a 2k x 2k Orius 830 CCD camera (Roper Scientific, Evry, France).

### Human tissue immunohistochemistry

Our study included specimens obtained from the brain collection “Hôpitaux Universitaires de l’Est Parisien – Neuropathologie du développement” (Biobank identification number BB-0033-00082). Informed consent was obtained for autopsy of the brain and histological examination. Our study included fetal brains obtained from spontaneous or medical abortion that did not display any significant brain pathology. After removal, brains were fixed with formalin for 5–12 weeks. Macroscopic analysis was performed to select samples that where embedded in paraffin, sliced in 7 μm sections and stained with hematein for a first histological analysis. Immunohistochemical analyses were performed on coronal slices that included the temporal telencephalic parenchyma and hippocampus. They were dewaxed and rinsed before incubation in citrate buffer (pH 9.0). Expression of MLC1 on the sections was detected using the Bond Polymer Refine Detection Kit (Leica) with specific antibodies and an immunostaining system (Bond III, Leica). Images were acquired using a slide scanner (Lamina, Perkin Elmer). Staining was analyzed using QuPath [34]. A QuPath pixel classifier was trained to discriminate between DAB-positive spots and background areas. We selected a pixel classifier that used a random trees algorithm and four features: a Gaussian filter to select intensity, and the three structure tensor eigenvalues to select thin elongated objects. The classifier was trained on manually annotated MLC1 spots and the background area on one image per developmental stage. When the result was satisfactory, the pixel classifier was used to detect MLC1 in selected regions of interest.

### Statistics

For all variables, the normality of data distribution was probed with using the Shapiro-Wilk test before the appropriate statistical test was chosen. Test names and sample sizes are indicated in the figure legends. Detailed results are presented in the supplementary tables.

## Results

### The absence of MLC1 results in accumulation of fluid in the brain but does not alter blood-brain barrier (BBB) integrity or the organization of the endothelial network

Astrocytes influence several properties of endothelial cells, such as BBB integrity [18–21]. We recently demonstrated that the postnatal maturation of the MLC1/GlialCAM complex in PvAPs from P5 to P15 coincides with the progressive increase in endothelium-specific proteins that contribute to BBB integrity, such as the tight junction protein claudin5 and the endothelial luminal ATP-binding cassette (ABC) efflux transporter P-glycoprotein (P-gP) - suggesting that PvAPs and the BBB mature in parallel [16, 17]. We therefore looked at whether the absence of MLC1 in the *Mlc1* KO mouse influenced postnatal endothelial maturation. To that end, we used qPCRs to characterize the expression of *Abcb1* (encoding P-gP) and *Cldn5* (encoding claudin5) on P5 and P15 and at 2 months (P60) on mRNAs extracted from whole brain microvessels (MVs) purified from WT and *Mlc1* KO animals [22] **(Fig. 1a; Table S1)**. In the WT, expression of all the analyzed mRNAs on P5 and P15 confirmed the progressive postnatal molecular maturation of endothelial cells (**Fig. 1a**). There were no differences between *Mlc1* KO mice and WT mice in this respect (**Fig. 1a**). Consistently, the protein levels of P-gP and claudin5 in purified MVs (on Western blots) were also similar in *Mlc1* KO and WT mice at all stages (**Fig. 1b; Table S2**). We next assessed BBB integrity in two-month-old *Mlc1* KO and WT mice. We first observed that the apparent diffusion coefficient (ADC, measured using MRI) was higher in *Mlc1* KO mice (**Fig. 1c; Fig. S1; Table S3).** Furthermore, the volume of the ventricles and the brain estimated from the anatomical T2-weighted MRI acquisition was larger in *Mlc1* KO mice than in WT mice **(Fig. S1c)**, although the relative ventricular volumes did not differ (**Fig. S1d**). These results reflected the previously described presence of fluid in the parenchyma in *Mlc1* KO mice [9]. However, this fluid accumulation was not related to leakage of the BBB. Indeed, the vascular volume measured by *in situ* brain perfusion of [^14^C] sucrose (a marker of vascular space and integrity [24]) was the same in *Mlc1* KO and WT mice and so suggested that the BBB was not leaky (**Fig. 1d; Table S4**) - even in the shear stress conditions (increased hydrostatic pressure: 180 mmHg) produced by the addition of human serum albumin to the perfusate [23] (**Fig. 1d**). Lastly, we assessed the endothelium architecture by analyzing vessel length, branching and tortuosity in the whole cleared somatosensory cortex of 2-month-old WT and *Mlc1* KO immunolabeled for the endothelium-specific protein Pecam-1 (**Fig. 1e-h; Table S5**). There were no differences between WT and *Mlc1* KO mice with regard to these architectural parameters, either in the parenchymal (**Fig. 1e, f)** or pial vasculature at the cortical surface (**Fig. 1g, h)**.

**Fig. 1.**
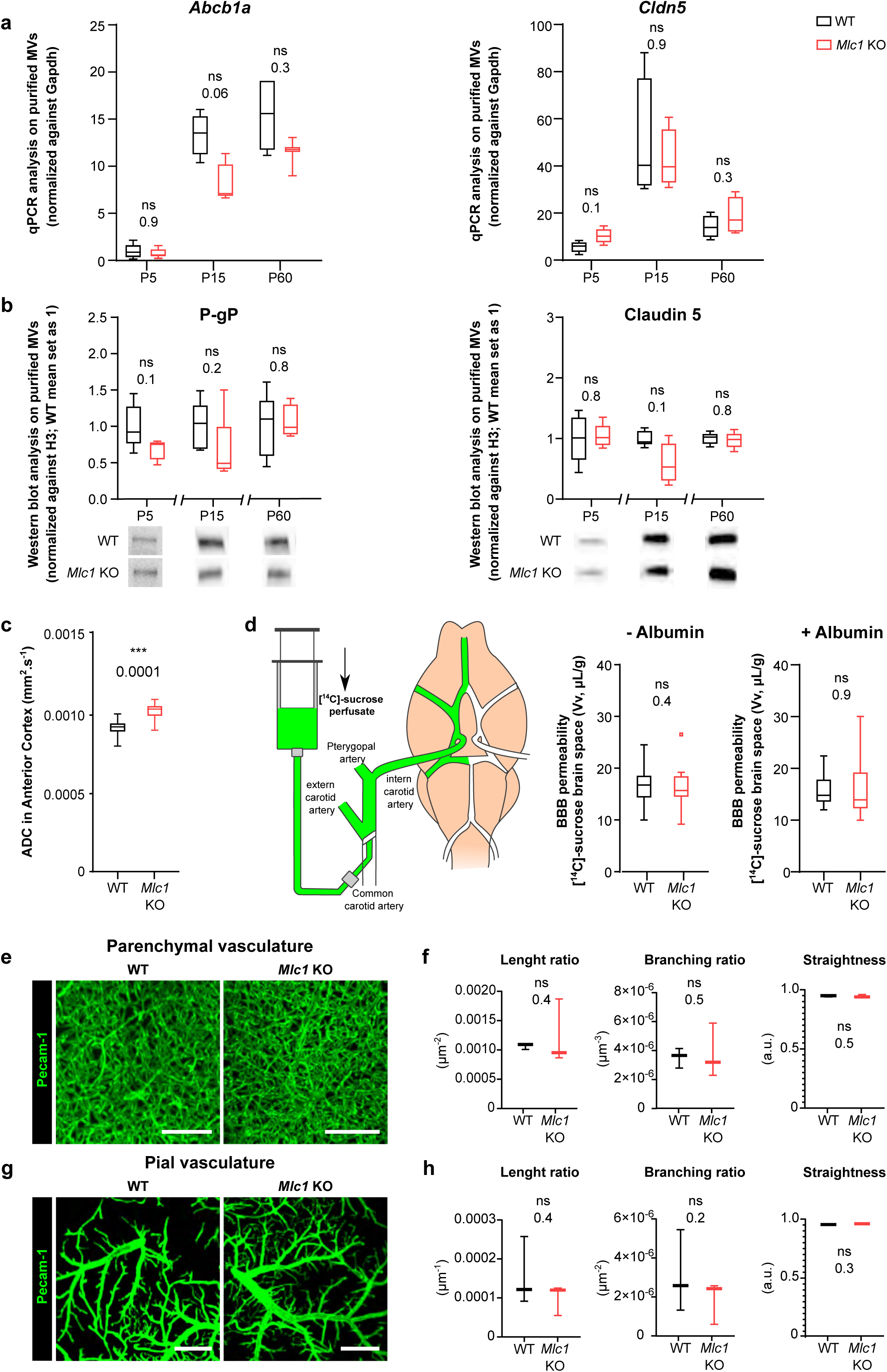
The absence of MLC1 has no effect on BBB integrity or the organization of the endothelial network. **a** mRNA expression (using qPCR) of *Abcb1* (encoding P-gP) and *Cldn5* (encoding claudin5) in MVs purified from WT and *Mlc1* KO whole brains on P5 and P15 and at 2 months (P60). Signals were normalized against *Gapdh*. Groups were compared using a two-tailed Mann-Whitney test. The data are presented as a Tukey box plot (n = 3 or 4 samples per genotype; mice per sample: 5 for P5; 3 for P15; 2 for P60)). Data are given in **Table S1**. **b** Western blot detection and analysis of P-gP and claudin5 in protein extracts from MVs purified from WT and *Mlc1* KO whole brains on P5 and P15 and at 2 months (P60). Signals were normalized against histone H3. Two-tailed Mann-Whitney test. The data are represented in a Tukey box plot (n = 4 or 5 sample per genotype; mice per sample: 5 for P5; 3 for P15; 2 for P60). Data are given in **Table S2**. **c** Apparent diffusion coefficient (ADC) values in the cortex of 2-month-old WT and *Mlc1* KO mice. Two-tailed Student’s T test. The data are represented in a Tukey box plot (n = 7 mice per genotype). Data are given in **Table S3**. **d** BBB integrity, assessed by measuring the brain vascular volume (Vv, in µL/g) after *in situ* brain perfusion with [^14^C]-sucrose and a normal hydrostatic vascular pressure (Albumin -; 120 mmHg) or an elevated hydrostatic vascular pressure (Albumin +; 180 mmHg) in 2-month-old WT (black) and *Mlc1* KO mice (red). Two-tailed Mann-Whitney test. The data are represented in a Tukey box plot (n = 8 WT and 9 *Mlc1* KO mice for Albumin -; n = 11 WT and 12 *Mlc1* KO mice for Albumin +). Data are given in **Table S4**. **e g** Representative 3D images of the endothelial architecture in cleared somatosensory cortex. Parenchymal (Z stack 320µm; scale bar: 100µm) **(e)** and pial **(g)** vessels (Z stack 50 µm; scale bar: 500µm) samples from 2-month-old WT and *Mlc1* KO mice, after immunolabeling for Pecam1. **f h** A comparative analysis of vessel length, branching and tortuosity in WT mice (black) and *Mlc1* KO mice (red) in the parenchymal cortex, normalized on sample volume **(f)** and cortical surface, normalized on sample surface **(h)**. One-tailed Mann-Whitney test. The data are represented in a Tukey box plot (n = 3 mice per genotype). Data are given in **Table S5**. *, p ≤ 0.05, **, p ≤ 0.01, ***, p ≤ 0.001, and ns: not significant.

Hence, the absence of MLC1 leads to fluid accumulation in the brain but has no obvious impact on the postnatal molecular maturation of ECs, BBB integrity, or endothelium architecture.

### MLC1 is crucial for contractile maturation of VSMCs, arterial perfusion, and neurovascular coupling

We recently showed that VSMCs mature postnatally, with the progressive acquisition of contractility from P5 onwards and the extension of the VSMC network [17]. Here, we investigated the VSMCs’ status in *Mlc1* KO mice during postnatal development. We first used qPCRs to compare the mRNA expression of *Acta2* (encoding smooth muscle actin, SMA) in MVs purified from WT and *Mlc1* KO whole brain on P5 and P15 and at 2 months [22] **(Fig. 2a; Table S1).** We also measured the mRNA expression of *Atp1b1* (a VSMC-specific gene stably expressed during postnatal development) as a marker of VSMC density in purified MVs [35, 36]. In WT mice, the level of contractile-protein-encoding mRNAs rose progressively from P5 to P15, while the level of *Atp1b1* mRNA remained stable **(Fig. 2a)**. In *Mlc1* KO mice, however, *Acta2* was significantly downregulated between on P5 and P15, and *Atp1b1* levels were unchanged **(Fig. 2a)**. We next analyzed the protein levels of SMA in MVs purified from WT and *Mlc1* KO whole brain on P5 and P15 and at 2 months by Western blot (**Fig. 2b; Table S2**). A small decrease in SMA expression was found in *Mlc1* KO MVs from P15 onwards although it became significant only at 2 months. Moreover, at 2 months, additional bands of lower molecular weights (resembling a degradation pattern) were detected **(Fig. 2b)**. To determine whether the decrease in SMA expression and the putative increase in its degradation were related to VSMC degeneration, we used an immunofluorescence assay to detect SMA on 2-month-old WT and *Mlc1* KO whole cleared somatosensory cortices (**Fig. 2c-f**). No discontinuities in the labeling were detected in the parenchymal (**Fig. 2c, d**) or pial vasculature (at the cortical surface) (**Fig. 2e, f**). Moreover, the SMA-positive vessels’ length, branching, tortuosity and number of anastomoses (analyzed only in pial vessels) were the same in *Mlc1* KO and WT mice (**Fig. 2c-f**). These findings indicate that the absence of MLC1 perturbs the developmental expression of SMA but does not affect the development of the vascular VSMC network.

**Fig. 2.**
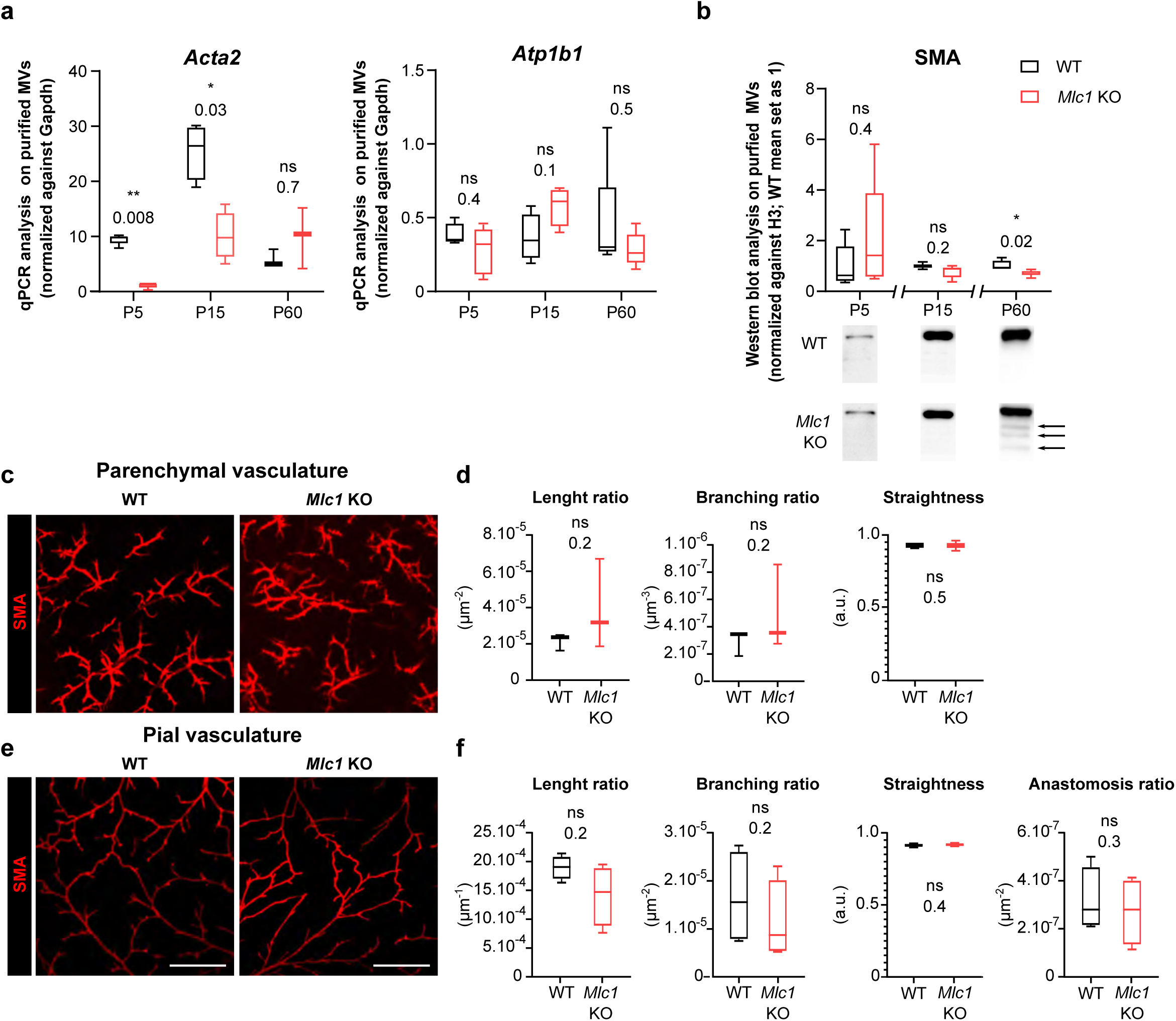
MLC1 is crucial for the molecular maturation of VSMC contractility. **a** qPCR results for *Acta2* (encoding SMA) and *Atp1b1* in MVs purified from WT and *Mlc1* KO whole brains on P5 and P15 and at 2 months. Signals are normalized against *Gapdh*. Two-tailed Mann-Whitney test. The data are represented in a Tukey box plot (n = 3 to 5 samples per genotype; mice per sample: 5 for P5; 3 for P15; 2 for P60)). Data are given in **Table S1**. **b** Western blot detection and analysis of SMA in protein extracts from MVs purified from WT and *Mlc1* KO whole brains on P5 and P15 and at 2 months (P60). Arrows indicate abnormally low SMA-positive bands. Signals were normalized against histone H3. Two-tailed Mann-Whitney test. The data are represented in a Tukey box plot (n = 4 or 5 samples per genotype; mice per sample: 5 for P5; 3 for P15; 2 for P60)). Data are given in **Table S2**. **c e** Representative 3D images of the VSMC arterial network in cleared somatosensory cortex. Parenchymal (Z stack 600 µm; scale bar: 100µm) **(c)** and pial vessels (Z stack 50 µm; scale bar: 500µm) **(e)** samples in 2-month-old WT and *Mlc1* KO mice after immunolabeling for SMA. **d f** Comparative analysis of arterial length, branching, tortuosity, and anastomosis in WT mice (black boxes) and *Mlc1* KO mice (red boxes) in the cortical parenchyma, normalized on sample volume (**d**) and at the cortical surface, normalized on sample surface (**f**). One-tailed Mann-Whitney test. The data are represented in a Tukey box plot (n = 3 mice per genotype). Data are given in **Table S5**. *, p ≤ 0.05, **, p ≤ 0.01, ***, p ≤ 0.001, and ns: not significant.

To further assess the functional consequences of this molecular change, we compared the *ex vivo* contractility of VSMCs in brain slices obtained from *Mlc1* KO and WT mice on P5 and P15 and at 2 months (**Fig. 3a-c; Table S6**). We recorded the vasomotor changes in cortical arterioles upon exposure for 2 min to a thromboxane A_2_ receptor agonist U46619 (9,11-dideoxy-11a,9a-epoxymethanoprostaglandin F2α, 5 nM), which acts directly on VSMCs to induce a reversible vasoconstriction. On P5, application of U46619 had a small effect on vessel diameter in both WT and *Mlc1* KO mice **(Fig. 3b, c)**. In contrast, a clear vasoconstriction was observed from P15 **(Fig. 3b, c)**. Strikingly, the amplitude and speed of vasoconstriction were significantly lower in *Mlc1* KO mice from P15 onwards (**Fig. 3b, c; Table S6**). These results indicate that the postnatal acquisition of contractility is impaired in the absence of MLC1.

**Fig. 3.**
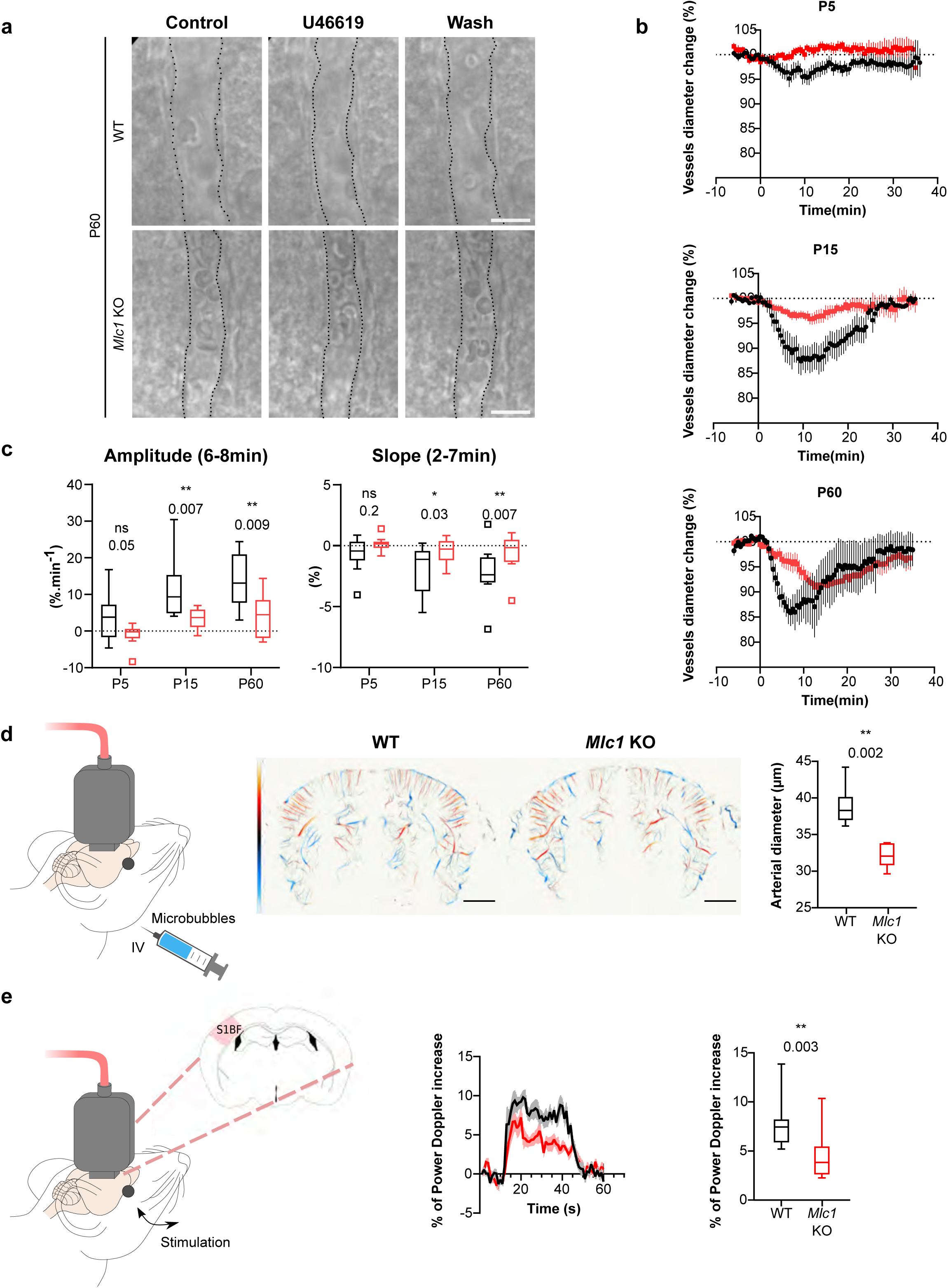
MLC1 is crucial for the postnatal acquisition of VSMC contractility, arterial diameter, and neurovascular coupling. **a-c** *Ex vivo* analysis of mean vascular constriction and dilation upon application of U46619 (50 nM) to somatosensory cortical slices from P5, P15 and 2-month-old WT mice (black traces) and *Mlc1* KO mice (red traces). **a** Representative infrared images of a cortical penetrating arteriole constriction in response to bath application of U46619 and dilation upon washing at P60. The vessel lumen is indicated by a dotted line. Scale bar: 10µm. **b** Contraction and dilation slopes on P5, P15 and P60. 0 min corresponds to the addition of U46619 to the recording chamber medium. The data are quoted as the mean ± SEM. **c** Analysis of the amplitude and slope of the contraction. Two-tailed Mann-Whitney test. The data are represented in a Tukey box plot (n = 13 vessels from WT and 13 *Mlc1* KO mice on P5; n = 9 vessels from WT and 12 *Mlc1* KO mice at P15; n = 8 vessels from WT and 11 *Mlc1* KO mice at P60; 3 mice per group). Data are given in **Table S6**. **d** *In vivo* ULM measurement of cortical arterial vessel diameter after intravenous microbubble injection in 2-month-old WT and *Mlc1* KO mice. **Left:** Schematic representation of the experiment; **Middle:** Cerebrovascular maps of WT and *Mlc1* KO mice, showing the arterial (red) and venous (blue) velocities in mm/s (scale bar: 0.15 cm); **Right:** Measurement of the penetrating arteries’ diameter using ULM imaging of an injected microbubbles. The data are represented in a Tukey box plot. Two-tailed Mann–Whitney test (n=6 WT mice and *Mlc1* KO mice). Data are given in **Table S7**. **e** *In vivo* fUS analysis of CBF in the somatosensory cortex after whisker stimulation. **Left:** Schematic representation of the experiment (S1bf: bregma −1.5 mm, somatosensory barrel field cortex); **Middle:** fUS power doppler signal traces during whisker stimulation of 2-month-old WT mice (black) and *Mlc1* KO mice (red) mice; **Right:** Quantification of the normalized CBF variations. Two-tailed Mann–Whitney test. Curves in transparency correspond to the SEM (n=11 WT mice and 12 *Mlc1* KO mice). Data are given in **Table S7**. *, p ≤ 0.05, **, p ≤ 0.01, ***, p ≤ 0.001, and ns: not significant.

Given this phenotype, we next hypothesized that arterial tonicity might be impaired in *Mlc1* KO mice. We first addressed this question by performing Ultrasound localization microscopy (ULM) *in vivo* imaging to reveal the brain vasculature at a microscopic resolution after intravenous microbubble injection (**Fig. 3d)**. *Mlc1* KO mice displayed significantly lower blood perfusion, which was suggestive of narrower penetrating arteries **(Fig. 3d; Table S7)**. Vasomotricity and cerebral blood flow (CBF) are tightly coupled to neuronal energy demand, in a process referred to as neurovascular coupling or functional hyperemia [37]. We then used functional Ultrasound (fUS) imaging to measure variations in CBF in the barrel cortex in response to whisker stimulation in 2-month-old WT and *Mlc1* KO mice [27, 38, 39] **(Fig. 3e; Table S7)**. The increase in CBF evoked by whisker stimulation was significantly smaller in *Mlc1* KO mice than in WT mice, indicating that neurovascular coupling was impaired in the KO mice **(Fig. 3e; Table S7)**.

In conclusion, the absence of MLC1 impairs the postnatal acquisition of contractile properties by VSMCs and impedes blood perfusion, vessel diameter and neurovascular coupling.

### The absence of MLC1 alters the PvAPs’ molecular maturation and perivascular cohesiveness

We next characterized the PvAPs in *Mlc1* KO mice by focusing on membrane proteins known to be strongly expressed in these structures [18]: the water channel aquaporin 4 (Aqp4), the adhesion protein GlialCAM, and the gap junction protein connexin 43 (Cx43). We first used immunofluorescence to analyze the proteins’ localization in PvAPs on brain sections on P5, P15 and at 2 months (**Fig. 4a**). Vessels were counterstained with isolectin B4. Interestingly, perivascular Aqp4, GlialCAM and Cx43 immunolabeling was almost undetectable (relative to WT) in *Mlc1* KO mice on P5 (**Fig. 4a**). In contrast, all proteins were detected normally in PvAPs from P15 onwards - with the exception of GlialCAM, which was not detected at any stage (as described previously [5]) (**Fig. 4a**). To further confirm and quantify these results, we assessed the levels of Cx43, GlialCAM and Aqp4 on Western blots of whole brain protein extracts on P5 and P15 and at 2 months (**Fig. 4b; Table S2**). In *Mlc1* KO mice, lower levels of all these proteins were observed but only on P5 (**Fig. 4b**). These data suggested that the molecular maturation of PvAPs is delayed in *Mlc1* KO mice.

**Fig. 4.**
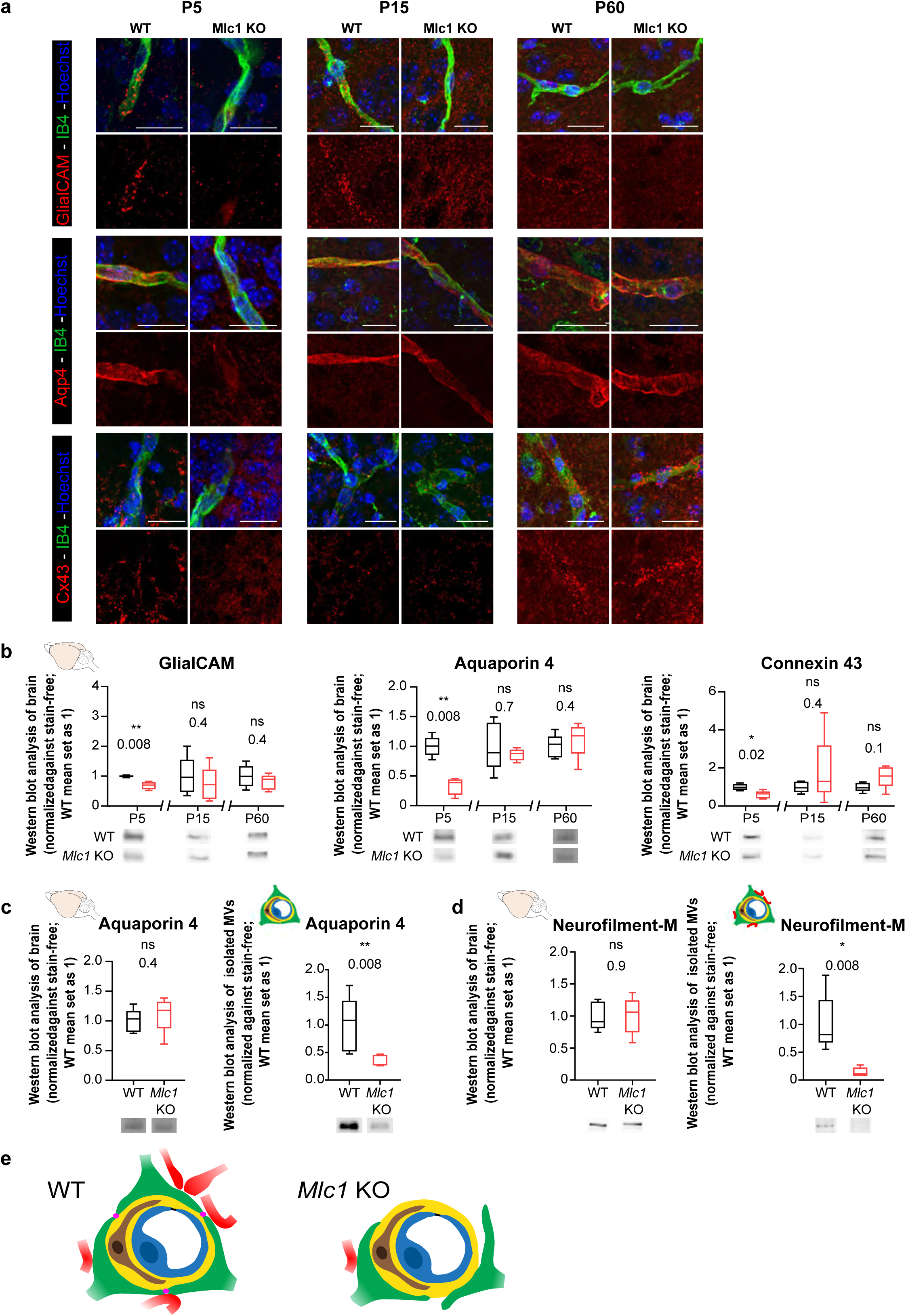
The absence of MLC1 alters the molecular maturation and adherence of PvAPs. **a** Representative confocal projection images of the immunofluorescent detection of GlialCAM, Aqp4 and Cx43 (red) on brain cortex sections from WT and *Mlc1* KO mice on P5, P15 and at 2 months. Vessels are stained with isolectin B4 (green) and nuclei are stained with Hoechst dye (blue). Scale bar: 20 µm. **b** Western blot detection and analysis of GlialCAM, Aqp4 and Cx43 in protein extracts from whole brains from WT and *Mlc1* KO mice on P5 and P15 and 2 months. **c** Western blot detection and analysis of Aqp4 in protein extracts from MVs purified from WT and *Mlc1* KO whole brains at 2 months (P60). **d** Western blot detection and analysis of NF-M in protein extracts from MVs purified from WT and *Mlc1* KO whole brains at 2 months (P60). Signals were normalized against stain-free membranes. Two-tailed Mann-Whitney test. The data are represented in a Tukey box plot (for whole brain: n = 5 samples per genotype (one mouse per sample); for purified MVs: n = 5 samples per genotype (2 mice per sample). Data are given in **Table S2**. **e** Schematic interpretation of the data showed in (**c**) and (**d**). In *Mlc1* KO mice, the PvAPs (green) and neuronal associated fibers (red) are lost during the MV purification process (BL, yellow; mural cell, brown; EC, blue). MLC1 is represented by pink dots in WT. *, p ≤ 0.05, **, p ≤ 0.01, ***, p ≤ 0.001, and ns: not significant.

We had demonstrated previously that PvAPs and the associated neuronal fibers remain attached to vessels during their mechanical purification (**Fig. 4e**) [22]. Surprisingly, Aqp4 levels were similar in whole brain extracts from 2-month-old WT and *Mlc1* KO mice but were significantly lower in extracts from *Mlc1* KO mechanically purified brain MVs (**Fig. 4c; Table S2**). The same was true for neurofilament protein M (NFM), a neuronal-specific intermediate filament protein present in neuronal fibers abutting PvAPs (**Fig. 4d**). NF-M was similarly present in whole brain extracts from WT and Mlc1 KO mice but was almost undetectable in 2-month-old *Mlc1* KO MVs (**Fig. 4d; Table S2**). These data suggested that the PvAPs and the associated neuronal fibers had detached from *Mlc1* KO brain vessels during the mechanical purification of MVs.

Taken as a whole, these results show that the absence of MLC1 delays the acquisition of Aqp4 and Cx43, disrupts (from P5 onwards) the membrane anchorage of GlialCAM, and reduces the cohesiveness of PvAPs at the vessel surface (**Fig. 4e**).

### The absence of MLC1 alters astrocyte morphology and polarity, PvAP morphology, and perivascular coverage

Our results suggested that in the absence of MLC1, PvAPs may not adhere properly to the vessel surface. Since adhesion, cell morphology and polarity are interdependent, we hypothesized that the absence of MLC1 could perturb the astrocytes’ morphology and polarity. We addressed this question by performing a Sholl analysis of glial fibrillary acid protein (GFAP)-immunolabeled astrocytic ramifications in the CA1 region of the hippocampus in 2-month-old mice (**Fig. 5a, b**). *Mlc1* KO astrocytes displayed a greater number of processes located between 15 to 25 µm from the soma (**Fig. 5b; Table S8**).

**Fig. 5.**
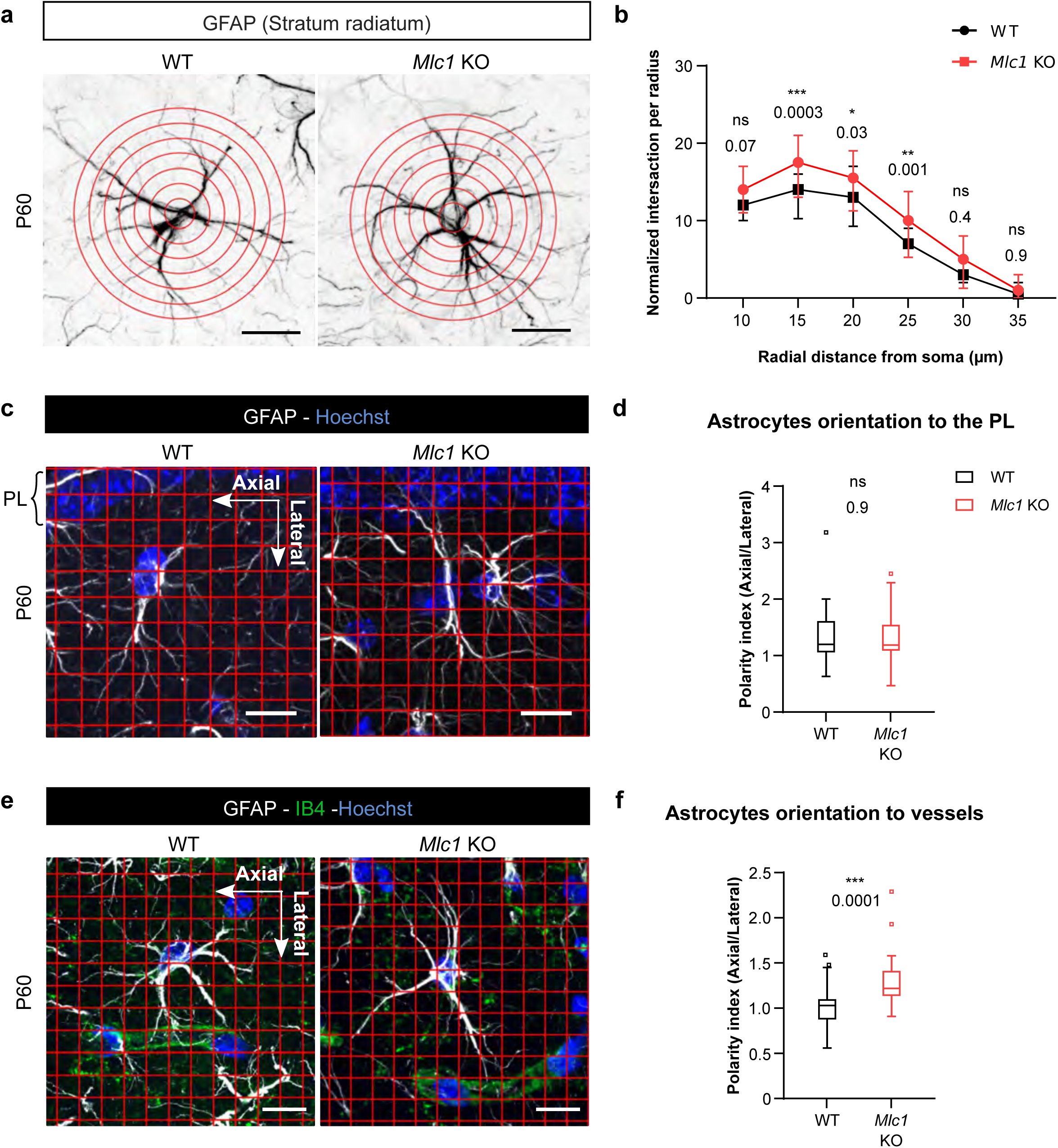
The absence of MLC1 alters astrocyte morphology and polarity. **a** A Sholl analysis of the ramification of hippocampal CA1 astrocytes immunolabeled for GFAP (black) in WT and *Mlc1* KO 2-month-old mice. Concentric circles are drawn starting from the astrocyte’s soma. Scale bar: 20µm. **b** Quantitative analysis of the astrocyte’s ramifications. Two-way ANOVA test followed by a Bonferroni *post hoc* test. Data are quoted as the median ± IQR (n=48 *Mlc1* KO cells; n=44 WT cells; n=3 mice per genotype). Data are given in **Table S8**. **c** Grid-baseline analysis of the orientation of the GFAP-immunolabeled astrocytic processes (white) toward the hippocampal pyramidal cell layer (PL) in WT and *Mlc1* KO 2-old-month mice. Nuclei are labeled with Hoechst dye (blue). Scale bar: 20µm. **d** Quantitative analysis of astrocyte process orientation toward the PL. The polarity index is the ratio between of axial GFAP contacts and lateral GFAP contacts. A polarity index of 1 means that there is no polarity. Two-tailed Mann-Whitney test. The data are represented in a Tukey box plot (n=52 *Mlc1* KO cells; n= 47 WT cells; 3 mice per genotype). Data are given in **Table S8**. **e** Grid-baseline analysis of the orientation of the GFAP immunolabeled astrocytic processes (white) toward vessels labeled with isolectin B4 (green) in WT and *Mlc1* KO 2-month-old mice. Nuclei are labeled with Hoechst dye (blue). Scale bar: 20µm. **f** Quantitative analysis of astrocyte process orientation toward vessels. The polarity index is the ratio between of axial GFAP contacts and lateral GFAP contacts. A polarity index of 1 means that there is no polarity. Two-tailed Mann-Whitney test. The data are represented in a Tukey box plot (n=41 *Mlc1* KO cells; n= 41 WT cells; 3 mice per genotype). Data are given in **Table S8**. *, p ≤ 0.05, **, p ≤ 0.01, ***, p ≤ 0.001, and ns: not significant.

In the hippocampal stratum radiatum, GFAP-positive astrocytic processes are normally polarized perpendicular to the pyramidal cell layer (PL) [40]. We evaluated this preferential orientation and measured the polarity index, which corresponds to the ratio between parallel (axial) and perpendicular (lateral) crossing points between GFAP-positive processes and a grid oriented with the PL (**Fig. 5c, d**). We found that both *Mlc1* KO and WT astrocytes were equally well oriented, with a polarity index > 1 (**Fig. 5d; Table S8**). However, when taking hippocampal vessels as the reference (**Fig. 5e, f)**, *Mlc1* KO astrocytes were abnormally oriented towards vessels in the axial plane (**Fig. 5e, f; Table S8).**

We next used transmission electron microscopy (TEM) to analyze the ultrastructural morphology of PvAPs in the cortex and hippocampus of 2-month-old WT and *Mlc1* KO mice, with a focus on vessels up to 10 μm in diameter (**Fig. 6)**. In WT mice, endothelial cells were joined by tight junctions (TJs) and were totally covered by PvAPs, which themselves were joined by gap junctions. Astrocytes and endothelial cells were separated by a thin, homogeneous, regular basal lamina (BL) (**Fig. 6a**). In *Mlc1* KO mice, the endothelium appeared to be unaltered: normal tight junctions (TJ) and BL, and no accumulation of intracellular vesicles (**Fig. 6b**). However, the astrocytes’ perivascular organization was drastically modified (**Fig. 6b-e**). We observed PvAPs surrounded by BL (**Fig. 6c**), or stacked on top of each other and joined by gigantic gap junction plaques, suggesting a loss of polarization (**Fig. 6d**). Some of the PvAPs interpenetrated each other (**Fig. S2b**). Astrocyte coverage was often discontinuous. In the free spaces, axons (recognizable by their microtubules) (**Fig. 6b, e**) and synapses (recognizable by the large number of vesicles in the presynaptic part and their electron dense postsynaptic density (**Fig. S2b**)) were found to be in direct contact with the endothelial BL. Accordingly, the percentage of MVs in which the endothelial BL was in direct contact with at least one neuronal process was higher in *Mlc1* KO mice than in WT mice (**Fig. 6f**). Despite the presence of these anomalies, no swelling was observed in *Mlc1* KO PvAPs vs. WT PvAPs (**Fig. 6b-e, g; Table S9)**.

**Fig. 6.**
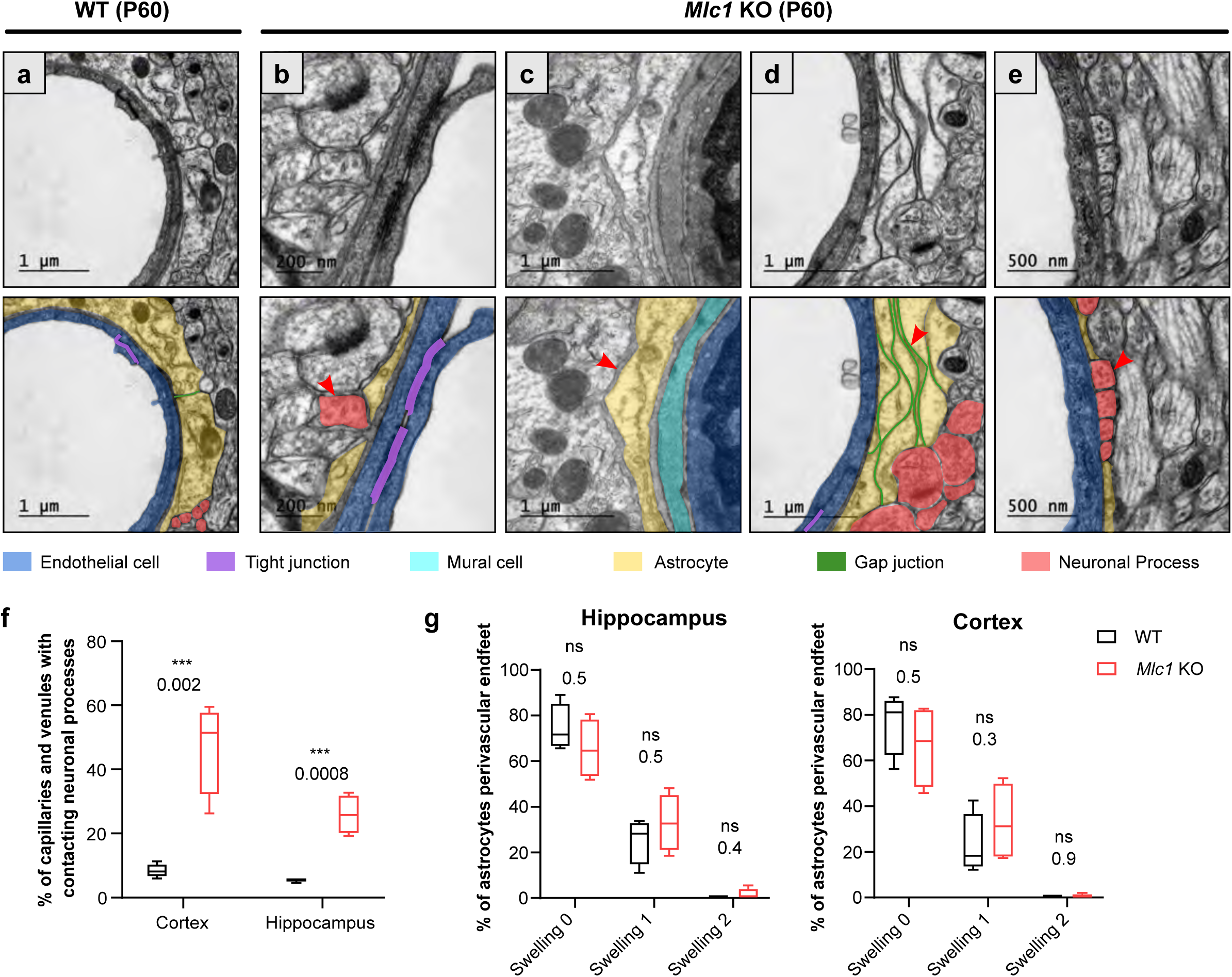
The absence of MLC1 alters PvAP polarity and perivascular coverage. **a-e** Representative TEM images of the GVU in the hippocampus of 2-month-old WT and *Mlc1* KO mice (n=3 mice per genotype). Images are presented in pairs, with artificial colors in the lower panel: PvAPs in yellow; gap junctions in green; axons or synapses in red; mural cell in light blue; endothelial cells in dark blue; and TJs in purple. **a** In WT mice, PvAPs fully cover endothelial cells linked by a TJ and surrounded by a continuous basal lamina (BL). **b-e.** Data from *Mlc1* KO mice. **b** PvAPs are separated by an axon which contacts the endothelial BL (arrow head). **c** A PvAP is surrounded by the BL (arrow head). **d** Several PvAPs are stacked on the top of each other and are linked by extended gap junctions (arrow head). **e** PvAPs are separated by 4 axons that directly contact the vascular BL (arrow head). **f** Quantification of capillaries and venules contacted by neural processes (axons or synapses). T-test. The data are represented in a Tukey box plot (n=399 *Mlc1* KO cortical vessels; n=301 *Mlc1* KO hippocampal vessels; n=286 WT cortical vessels; n=287 WT hippocampal vessels; n=3 mice per genotype). **g** Percentage of vessels contacted by a normal PvAP (swelling 0), moderately swollen PvAP (swelling 1), or edematous PvAP (swelling 2) in hippocampus and cortex of 2-month-old mice. Two-tailed Mann-Whitney test. The data are represented in a Tukey box plot (n=399 *Mlc1* KO cortical vessels; n=301 *Mlc1* KO hippocampal vessels; n=286 WT cortical vessels; n=287 WT hippocampal vessels; n=3 mice per genotype). Data for (**f)** and (**g)** are given in **Table S9**. *, p ≤ 0.05, **, p ≤ 0.01, ***, p ≤ 0.001, and ns: not significant.

Taken as a whole, these results indicate that MLC1 is required for the astrocytes’ morphology and polarity. Within the GVU, the absence of MLC1 greatly alters PvAPs’ polarization, morphology and perivascular coverage.

### The absence of MLC1 modifies the parenchymal circulation of cerebrospinal fluid

Several groups have reported a causal link between PvAP disorganization and impaired parenchymal CSF transport [41, 42]. We tested this hypothesis by observing (using T1-weighted MRI) the parenchymal distribution of DOTA-Gadolinium (Gd) injected in CSF through the cisterna magna in 2-month-old mice (**Fig. 7a**). Contrast-enhanced T1 mapping was used to quantify changes over time in the concentration of DOTA-Gd within the brain tissue. In line with the conventional models of CSF solute circulation, tracers injected into the cisterna magna dispersed into the arachnoid space and then entered the parenchyma through the perivascular spaces (**Fig. 7b**). As expected, a high concentration of DOTA-Gd was detected at the border of the brain as soon as 10 minutes after injection in both WT and *Mlc1* KO mice; this reflected the initial dispersion of contrast within the subarachnoid space. Forty-five minutes after injection, the brain tissue concentration rose as the DOTA-Gd penetrated into the parenchyma through the perivascular spaces (**Fig. 7c**). For both genotypes, the distribution of DOTA-Gd appeared to be in line with previous observations [43]. However, examination of the quantitative maps suggested that DOTA-Gd transport in the CSF was lower in *Mlc1* KO mice than in WT mice. We analyzed the time courses of DOTA-Gd dispersion in the cerebellum, midbrain, septal area, and cortex (**Fig. 7d-g**). As compared to the WT mice, the mean DOTA-Gd concentration was lower in the midbrain (**Fig. 7e**) and the slope was lower in the cerebellum, midbrain, and cortex in the *Mlc1* KO mice (**Fig. 7d, e, g**).

**Fig 7.**
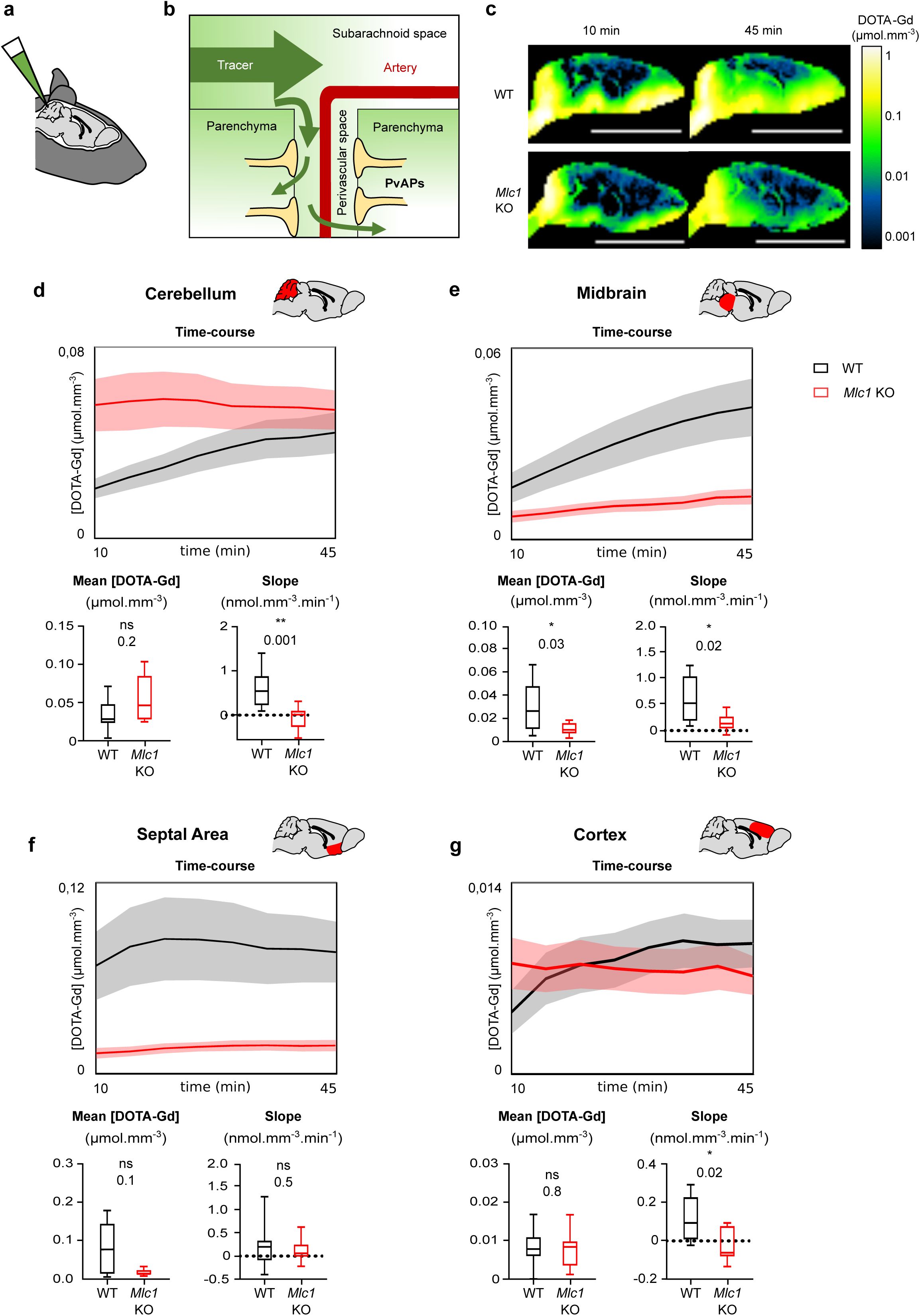
Contrast-enhanced MRI reveals reduced tracer dispersion from the CSF to the parenchyma in *Mlc1* KO mice. 1 µL DOTA-Gd was injected into the mouse’s CSF through the cisterna magna (**a**). After injection, the tracer disperses through the brain into the subarachnoid space before entering the deep parenchyma through the perivascular spaces (**b**). Quantitative contrast maps in WT and *Mlc1* KO mice, 10 and 45 minutes after contrast injection (scale bar: 1cm) (**c**). Based on the dynamic acquisitions, the changes over time in contrast agent concentration were extracted, and the mean contrast concentration and the contrast slope were calculated for the cerebellum, midbrain, septal area, and cortex. Two-tailed Mann-Whitney test. The data are represented in a Tukey box plot (n = 9 per genotype (8 in the cortex)). Data are given in **Table S3**. *, p ≤ 0.05, **, p ≤ 0.01, ***, p ≤ 0.001, and ns: not significant.

Thus, the absence of MLC1 impairs intraparenchymal circulation and clearance of CSF.

### The absence of MLC1 impacts the development of astrocyte morphology, polarity and perivascular coverage

MLC1 expression is a progressive process that starts in the mouse at P5 and completes at P15 [16]. In contrast, astrocyte ramification in the *stratum radiatum* increases greatly between P8 and P16 [40]. We therefore wondered whether the morphological and polarity defects observed in adult *Mlc1* KO astrocytes might be caused by an impairment development. Hence, we analyzed the morphology and orientation of GFAP-immunolabeled astrocytic processes on P10 and P15 [40] **(Fig. 8; Table S8)**. As observed at 2 months, *Mlc1* KO astrocytes had a larger number of processes at 10 to 25 µm from the soma (**Fig. 8a-d; Table S8**) and an abnormal axial orientation towards the hippocampal vessels on P10 and P15, relative to WT astrocytes **(Fig. 8 e-h; Table S8)**.

**Fig. 8.**
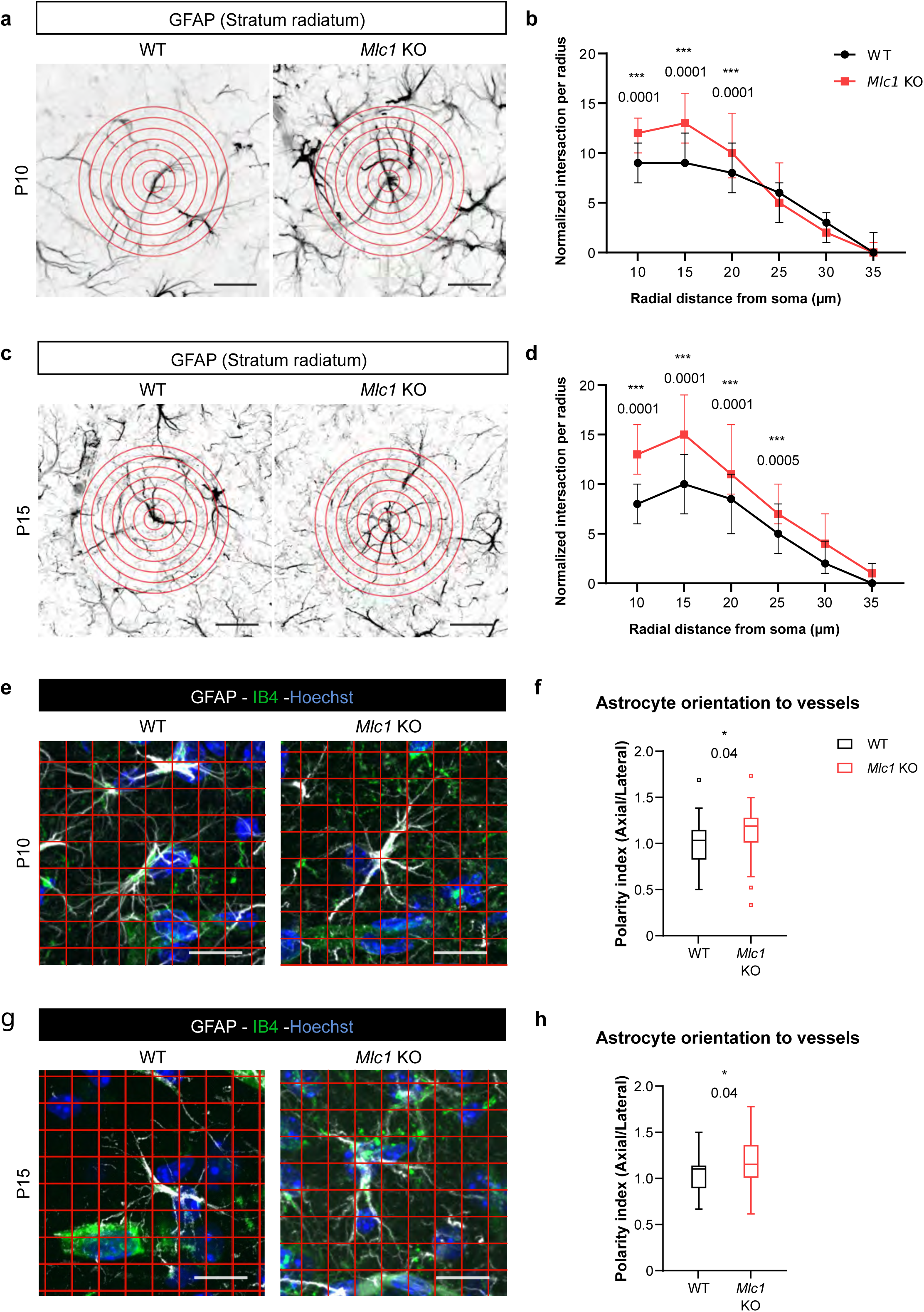
The absence of MLC1 alters the postnatal development of astrocyte morphology and polarity. **a-d.** A Sholl analysis of astrocyte ramification in WT and *Mlc1* KO mice on P10 (**a b**) and P15 (**c b**). Concentric circles are drawn, starting from the astrocyte’s soma. Scale bar: 20µm. **b d** Quantitative analyses of astrocyte ramification on P10 (**a**) and P15 (**c**). Two-way ANOVA test followed by a Bonferroni *post hoc* test (P10: n=53 *Mlc1* KO cells; n=49 WT cells; n=3 mice per genotype; P15: n=51 *Mlc1* KO cells; n=50 WT cells; n=3 mice per genotype). Data are presented in **Table S8**. **e-h** Grid analysis of astrocyte polarity towards vessel in WT and *Mlc1* KO on P10 (**e**) and P15 (**g**). Scale bar: 20µm. **f-h** Quantitative analysis of astrocyte process orientation toward vessels on P10 (**f**) and P15 (**g**). The polarity index is the ratio between of axial GFAP contacts and lateral GFAP contacts. A polarity index of 1 means that there is no polarity. Two-tailed Mann-Whitney test. The data are represented in a Tukey box plot (P10: n=37 *Mlc1* KO cells; n=26 WT cells; n=3 mice per genotype; P15: n=48 *Mlc1* KO cells; n=46 WT cells; n=3 mice per genotype). Data are presented in **Table S8**. *, p ≤ 0.05, **, p ≤ 0.01, ***, p ≤ 0.001, and ns: not significant.

The time course of perivascular astrocyte coverage has not previously been described. Here, we used TEM to quantify the percentage of the perivascular diameter covered by PvAPs in *Mlc1* KO and WT cortex and hippocampus on P5, P10, P15 and at 2-months (**Fig. 9; Table S9**). On P5, the PvAPs covered about half of the vessel’s circumference, and the WT and *Mlc1* KO samples did not differ significantly in this respect (**Fig. 9b**). The perivascular astrocyte coverage increased dramatically between P5 and P15 and was almost complete on P15 in WT mice (**Fig. 9b**). In great contrast, however, *Mlc1* KO mice displayed a lower percentage of PvAP coverage from P10 onwards, and a large number of neuronal processes were inserted into the uncovered vascular areas (**Fig. 9a; Table S9)**.

**Fig. 9.**
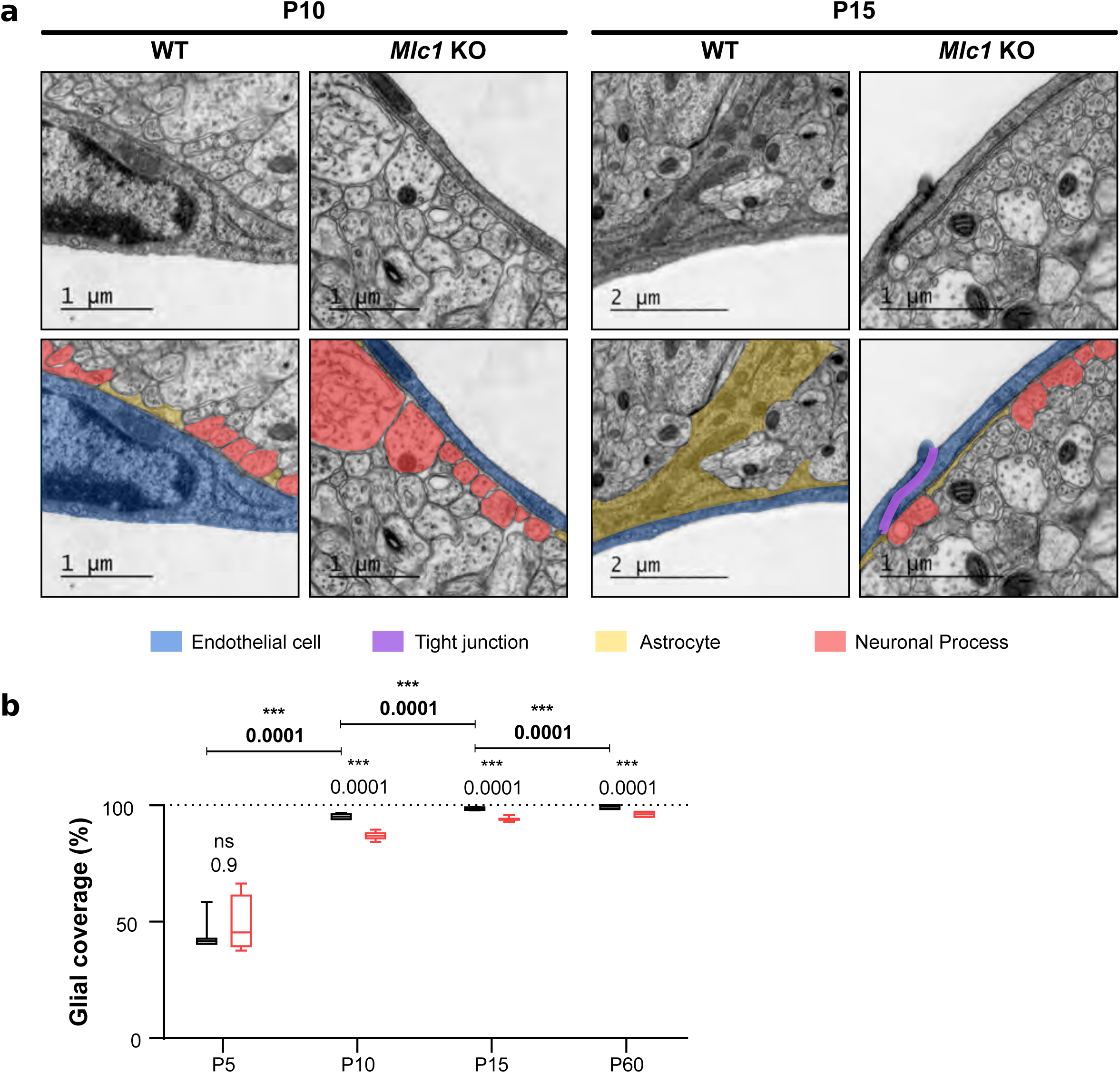
The absence of MLC1 alters the postnatal development of astrocyte perivascular coverage. **a** Representative TEM images of the gliovascular interface in the cortex of WT and *Mlc1* KO mice on P10 and P15. Images are presented in pairs, with artificial colors in the lower panel: PvAPs in yellow; axons in red; TJs in purple; and endothelial cells in blue. **b** Percentage of the vessel diameter covered by PvAPs in the cortex of WT and *Mlc*1 KO mice on P5, P10, P15, and P60. Two-tailed Mann-Whitney test. The data are represented in a Tukey box plot (n = 46 vessels from WT mice and 68 *Mlc1* KO mice on P5, n=4 mice per genotype; n=121 vessels from WT mice and 81 *Mlc1* KO mice on P10, n=3 mice per genotype; n=207 vessels from WT mice and 144 *Mlc1* KO mice at P15, n=4 mice per genotype; n=143 vessels from WT mice and 134 *Mlc1* KO mice at P60, n=3 mice per genotype). Data are presented in **Table S9**. *, p ≤ 0.05, **, p ≤ 0.01, ***, p ≤ 0.001, and ns: not significant.

MLC1 is therefore required for the development of astrocyte morphology and polarity. Our results demonstrated for the first time that the perivascular astrocyte coverage increases rapidly between P5 and P15. This process is impaired in *Mlc1* KO mice and results in incomplete perivascular PvAP coverage and direct contact between infiltrating neuronal processes and the endothelial basal lamina; the processes normally remain behind the perivascular astrocyte layer. MLC1 is therefore critical for the normal postnatal development of perivascular astrocyte coverage.

## Discussion

*MLC1* (the main gene involved in MLC) encodes an astrocyte-specific protein located in PvAPs, where it forms a junctional complex with GlialCAM. Our previous research showed that in the mouse, this complex forms progressively after birth (between P5 and P15) [16]. Our present work demonstrated that the P5-P15 time window is also important for the formation of perivascular astrocyte coverage. On P5, PvAPs covered only 50% of the vascular surface, and this coverage increased rapidly until completion on P15. The absence of MLC1 delays the PvAPs’ molecular maturation, with a transient decrease in Aqp4 and Cx43 protein levels on P5. GlialCAM (whose anchorage in PvAP membranes depends on MLC1) is distributed diffusely, as described previously [5]. In the absence of MLC1, impairments in the PvAPs are observed from P10 onwards, with incomplete perivascular coverage and direct contact between neuronal components and the vessel wall. Remodeling of the gliovascular interface is accompanied by a loss of astrocyte polarity. Firstly, the astrocytes have a greater number of ramifications, which tend to project towards the blood vessels. Secondly, there are changes in the stacking and interpenetration of PvAPs, the presence of BL on their parenchymal side, and the loss of perivascular cohesiveness. Taken as a whole, these data demonstrate that MLC1 is crucial for the development of astrocyte polarity and perivascular coverage.

What impact, then, does the lack of MLC1 have on the gliovascular physiology? The incomplete perivascular astrocyte coverage, the abnormal astrocytic polarity, and the loss of PvAP cohesiveness probably alter the “barrier” formed by PvAPs around the vessels. In turn, this might greatly affect perivascular homeostasis and astrocyte signaling towards the vascular compartment [19, 44]. The astrocytes’ perivascular organization is thought to be crucial for regulating the fluxes of CSF and interstitial fluid (ISF) into the parenchyma [45]. These alterations might therefore be directly linked to the loss of Gd drainage observed in *Mlc1* KO mice. Importantly, in the absence of BBB breakdown, impaired parenchymal circulation of CSF/ISF might (i) contribute to the fluid accumulation and megalencephaly observed in the *Mlc1* KO mouse model [9] and in patients with MLC [46], (ii) lead to the progressive accumulation of harmful molecules in the brain, and (iii) thus increase susceptibility to neural disorders [47].

Unlike skeletal or cardiac muscle, smooth muscle cells are not terminally differentiated; they are extremely plastic cells that are constantly integrating signals from their local environment and then expressing the appropriate patterns of genes. Smooth muscle cells undergo profound phenotypic changes in response to variations in the local environment [48, 49]. By perturbing perivascular homeostasis, the abnormal development of PvAPs in the absence of MLC1 might affect the postnatal acquisition of VSMCs’ contractile properties and thus result in hypoperfusion and defective neurovascular coupling. This hypothesis is supported by the fact that perivascular astrocyte coverage, MLC1 expression [16] and VSMC contractile differentiation [17] develop concomitantly. Interestingly, deletion of astrocytic laminin γ1 was shown to lead to the loss of VSMC contractile proteins [50], which indicated a functional link between astrocytes and VSMCs. Our data now suggest that astrocytes have a critical role in the postnatal differentiation of contractile VSMCs. Alteration of VSMC contractility might influence brain perfusion and neurovascular coupling, which are critical for oxygen and nutrient delivery to neurons [37]; this impairment might compromise neuronal and cerebral functions.

In *Mlc1* KO mice, myelin vacuolation has been shown to develop only from 3 months onwards [9, 5]. Thus, PvAP and VSMC maturation defects might be primary events in the pathogenesis of MLC. The resulting defects in perivascular homeostasis and neurovascular coupling might then contribute to progressive intramyelinic edema. In an attempt to move closer to the context of MLC in humans, we used immunohistochemical techniques to analyze the development of perivascular expression of MLC1 in human cortical sections from 15 weeks of gestation (wg) to 17 years of age (**Fig. S3**). Interestingly, perivascular MLC1 was detected as early as 15 wg and remained stable thereafter. These results suggested that the MLC1/GlialCAM complex and the astrocyte’s perivascular coverage are initiated prenatally, as already suggested by earlier observations of perivascular GFAP and AQP4 expression [51]. In the absence of MLC1, impaired PvAP formation might therefore occur *in utero* in individuals subsequently diagnosed with MLC. This impairment might deregulate perivascular homeostasis and then the postnatal differentiation of VSMC contractility.

In conclusion, we showed that the astrocyte-specific protein MLC1, which absence causes MLC, is critical for the postnatal development of perivascular astrocyte coverage, the acquisition of VSMC contractility, and parenchymal CFS/ISF efflux. Taken as a whole, these data indicate that MLC could be considered primarily as an early developmental disorder of the GVU. Moreover, our results shed light on the role of astrocytes in the postnatal the acquisition of VSMC contractility, a crucial component of neurovascular coupling in the brain. Our study illustrates how looking at physiopathological processes in a rare disease can enlighten important aspects of the brain’s physiology.

## Acknowledgements

We are grateful to the donors who support the charities and charitable foundations cited below. This work was funded by grants from the *Association Européenne contre les Leucodystrophies* (ELA, grant reference ELA2012-014C2B), the *Fondation pour la Recherche Médicale* (FRM, grant reference AJE20171039094) and the *Fondation Maladie Rares* (grant reference 20170603). A. Gilbert’s PhD was funded by the FRM (grant reference: PLP20170939025p60) and ELA (grant reference: ELA2012-014C2B). The creation of the Center for Interdisciplinary Research in Biology (CIRB) was funded by the “Fondation Bettencourt Schueller”. We thank Fawzi Boumezbeur, Aloïse Mabondzo, Corinne Blugeon, Laurent Jourdren and Stéphane Le Crom for helpful discussions. We thank Isabelle Bardou for her help writing ethical documents. We thank Louise Charpentier and Ines Masurel for their help analyzing astrocyte morphology and polarity. Lastly, we thank Virginie Mignon for help with the TEM analysis. Despite our efforts, our work has not received any funding from the French National Agency for Research (ANR).

## Supplementary information

**Fig. S1 The absence of MLC1 causes overall brain swelling**

Anatomical T2-weighted MRI **(a)**; wich representative images of WT and KO mice are shown **(b)**; was used to measure the brain volume **(c)** and the ventricles relative volume **(d)**. The ADC was calculated for the midbrain **(E)**, septal area **(F)**, and thalamus **(G)**. Two-tailed Student’s T test. The data are represented in a Tukey box plot (n=7 mice per genotype). Data are presented in **Table S3**. *, p ≤ 0.05, **, p ≤ 0.01, ***, p ≤ 0.001, and ns: not significant.

**Fig. S2 Examples of changes in the architecture of GVU in 2-month-old *Mlc1* KO mice**

**a-c** Representative TEM images of the GVU in the cortex of 2-month-old WT mice **(a)** and *Mlc1* KO mice **(b)** (n=3 mice per genotype). Images are presented in pairs, with artificial colors in the lower panel: PvAPs in yellow; gap junctions in green; synapse in orange; endothelial cells in dark blue; mural cell in light blue; and TJs in purple. **a** A WT sample showing continuous PvAP coverage around an endothelial cell. The PvAPs are linked by gap junctions **b** An *Mlc1* KO sample: **Left:** A synapse contacts the endothelial BL (arrowhead); **Middle:** A PvAP interdigitates into another PvAP and forms a large annular gap junction (arrowhead); **Right:** Stacked PvAPs surrounded by BL (arrowhead).

**Fig. S3: Developmental perivascular expression of MLC1 in the human cortex**

**a-e** Immunohistochemical analysis of MLC1 in the developing human cortex. **a-e** Representative images of MLC1-immunostained cortical slices (left) and a higher magnification image of the parenchyma in the boxed areas (right) at the prenatal stage (15 to 39 wg) **(a)**; 0 to 1 year of age (B); 3 to 4 years of age **(c)**; 10 to 13 years of age (D); 16 and 17 year of age **(e)**. Scale bar: 100µm. MLC1 immunostaining (arrowheads) was revealed with DAB. **f.** DAB intensity was quantified and quoted as the mean ± SD. We applied the Kruskal-Wallis test (overall, in bold) and a one-tailed Mann-Whitney test (for comparing stages). The number of samples per developmental age was 5 for prenatal, 5 for 0-1 years, 4 for 3-4 years, 4 for 10-13 years, and 2 for 16-17 years. Data are presented in

**Table S1 qPCRs data**

**Table S2 Western blot data**

**Table S3 MRI data**

**Table S4 in situ perfusion data**

**Table S5 Immunolabeling data**

**Table S6 Ex vivo VSMC contractility data**

**Table S7 ULM and fUS data**

**Table S8 Measurements of astrocyte morphology and polarity**

**Table S9 Electron microscopy data**

**Table S10 Immunohistochemistry data**

**Table S11 Resource table**

## References

1. Topcu M, Gartioux C, Ribierre F, Yalcinkaya C, Tokus E, Oztekin N, et al. Vacuoliting megalencephalic leukoencephalopathy with subcortical cysts, mapped to chromosome 22qtel. Am J Hum Genet. 2000;66(2):733–9.

2. Leegwater PA, Yuan BQ, van der Steen J, Mulders J, Konst AA, Boor PK, et al. Mutations of MLC1 (KIAA0027), encoding a putative membrane protein, cause megalencephalic leukoencephalopathy with subcortical cysts. Am J Hum Genet. 2001;68(4):831–8.

3. Duarri A, Teijido O, Lopez-Hernandez T, Scheper GC, Barriere H, Boor I, et al. Molecular pathogenesis of megalencephalic leukoencephalopathy with subcortical cysts: mutations in MLC1 cause folding defects. Hum Mol Genet. 2008;17(23):3728–39.

4. Lanciotti A, Brignone MS, Visentin S, De Nuccio C, Catacuzzeno L, Mallozzi C, et al. Megalencephalic leukoencephalopathy with subcortical cysts protein-1 regulates epidermal growth factor receptor signaling in astrocytes. Hum Mol Genet. 2016;25(8):1543–58.

5. Hoegg-Beiler MB, Sirisi S, Orozco IJ, Ferrer I, Hohensee S, Auberson M, et al. Disrupting MLC1 and GlialCAM and ClC-2 interactions in leukodystrophy entails glial chloride channel dysfunction. Nat Commun. 2014;5:3475.

6. Wang MX, Ray L, Tanaka KF, Iliff JJ, and Heys J. Varying perivascular astroglial endfoot dimensions along the vascular tree maintain perivascular-interstitial flux through the cortical mantle. Glia. 2020.

7. Ridder MC, Boor I, Lodder JC, Postma NL, Capdevila-Nortes X, Duarri A, et al. Megalencephalic leucoencephalopathy with cysts: defect in chloride currents and cell volume regulation. Brain. 2011;134(Pt 11):3342–54.

8. Capdevila-Nortes X, Lopez-Hernandez T, Apaja PM, Lopez de Heredia M, Sirisi S, Callejo G, et al. Insights into MLC pathogenesis: GlialCAM is an MLC1 chaperone required for proper activation of volume-regulated anion currents. Hum Mol Genet. 2013;22(21):4405–16.

9. Dubey M, Bugiani M, Ridder MC, Postma NL, Brouwers E, Polder E, et al. Mice with megalencephalic leukoencephalopathy with cysts: a developmental angle. Ann Neurol. 2015;77(1):114–31.

10. Bugiani M, Dubey M, Breur M, Postma NL, Dekker MP, Ter Braak T, et al. Megalencephalic leukoencephalopathy with cysts: the Glialcam-null mouse model. Ann Clin Transl Neurol. 2017;4(7):450–65.

11. Jeworutzki E, Lopez-Hernandez T, Capdevila-Nortes X, Sirisi S, Bengtsson L, Montolio M, et al. GlialCAM, a protein defective in a leukodystrophy, serves as a ClC-2 Cl(-) channel auxiliary subunit. Neuron. 2012;73(5):951–61.

12. Estevez R, Elorza-Vidal X, Gaitan-Penas H, Perez-Rius C, Armand-Ugon M, Alonso-Gardon M, et al. Megalencephalic leukoencephalopathy with subcortical cysts: A personal biochemical retrospective. Eur J Med Genet. 2018;61(1):50–60.

13. Benesova J, Rusnakova V, Honsa P, Pivonkova H, Dzamba D, Kubista M, et al. Distinct expression/function of potassium and chloride channels contributes to the diverse volume regulation in cortical astrocytes of GFAP/EGFP mice. PLoS One. 2012;7(1):e29725.

14. Elorza-Vidal X, Sirisi S, Gaitan-Penas H, Perez-Rius C, Alonso-Gardon M, Armand-Ugon M, et al. GlialCAM/MLC1 modulates LRRC8/VRAC currents in an indirect manner: Implications for megalencephalic leukoencephalopathy. Neurobiol Dis. 2018;119:88–99.

15. Gilbert A, Vidal XE, Estevez R, Cohen-Salmon M, and Boulay AC. Postnatal development of the astrocyte perivascular MLC1/GlialCAM complex defines a temporal window for the gliovascular unit maturation. Brain Struct Funct. 2019;224(3):1267–78.

16. Slaoui L, Gilbert A, Federici L, Rancillac A, Gelot A, Favier M, et al. In mice and humans, the brain’s blood vessels mature postnatally to acquire barrier and contractile properties. BioRXiv. 2021.

17. Cohen-Salmon M, Slaoui L, Mazare N, Gilbert A, Oudart M, Alvear-Perez R, et al. Astrocytes in the regulation of cerebrovascular functions. Glia. 2021;69(4):817–41.

18. Boulay AC, Saubamea B, Decleves X, and Cohen-Salmon M. Purification of Mouse Brain Vessels. J Vis Exp. 2015;105(105).

19. Dagenais C, Rousselle C, Pollack GM, and Scherrmann JM. Development of an in situ mouse brain perfusion model and its application to mdr1a P-glycoprotein-deficient mice. J Cereb Blood Flow Metab. 2000;20(2):381–6.

20. Ezan P, Andre P, Cisternino S, Saubamea B, Boulay AC, Doutremer S, et al. Deletion of astroglial connexins weakens the blood-brain barrier. J Cereb Blood Flow Metab. 2012;32(8):1457–67.

21. Iadecola C. The Neurovascular Unit Coming of Age: A Journey through Neurovascular Coupling in Health and Disease. Neuron. 2017;96(1):17–42.

22. Hingot V, Brodin C, Lebrun F, Heiles B, Chagnot A, Yetim M, et al. Early Ultrafast Ultrasound Imaging of Cerebral Perfusion correlates with Ischemic Stroke outcomes and responses to treatment in Mice. Theranostics. 2020;10(17):7480–91.

23. Osmanski BF, Pezet S, Ricobaraza A, Lenkei Z, and Tanter M. Functional ultrasound imaging of intrinsic connectivity in the living rat brain with high spatiotemporal resolution. Nat Commun. 2014;5:5023.

24. Bertolo A, Nouhoum M, Cazzanelli S, Ferrier J, Mariani JC, Kliewer A, et al. Whole-Brain 3D Activation and Functional Connectivity Mapping in Mice using Transcranial Functional Ultrasound Imaging. J Vis Exp. 2021(168).

25. Hwang J, Vu HM, Kim MS, and Lim HH. Plasma membrane localization of MLC1 regulates cellular morphology and motility. Mol Brain. 2019;12(1):116.

26. Nixdorf-Bergweiler BE, Albrecht D, and Heinemann U. Developmental changes in the number, size, and orientation of GFAP-positive cells in the CA1 region of rat hippocampus. Glia. 1994;12(3):180–95.

27. Kress BT, Iliff JJ, Xia M, Wang M, Wei HS, Zeppenfeld D, et al. Impairment of paravascular clearance pathways in the aging brain. Ann Neurol. 2014;76(6):845–61.

28. Haj-Yasein NN, Jensen V, Ostby I, Omholt SW, Voipio J, Kaila K, et al. Aquaporin-4 regulates extracellular space volume dynamics during high-frequency synaptic stimulation: a gene deletion study in mouse hippocampus. Glia. 2012;60(6):867–74.

29. Iliff JJ, Lee H, Yu M, Feng T, Logan J, Nedergaard M, et al. Brain-wide pathway for waste clearance captured by contrast-enhanced MRI. Journal of Clinical Investigation. 2013;123(3):1299–309.

30. Abbott NJ, Ronnback L, and Hansson E. Astrocyte-endothelial interactions at the blood-brain barrier. Nat Rev Neurosci. 2006;7(1):41–53.

31. Yao LL, Hu JX, Li Q, Lee D, Ren X, Zhang JS, et al. Astrocytic neogenin/netrin-1 pathway promotes blood vessel homeostasis and function in mouse cortex. J Clin Invest. 2020;130(12):6490–509.

32. Abbott NJ, Pizzo ME, Preston JE, Janigro D, and Thorne RG. The role of brain barriers in fluid movement in the CNS: is there a ‘glymphatic’ system? Acta Neuropathol. 2018;135(3):387–407.

33. van der Knaap MS, Boor I, and Estevez R. Megalencephalic leukoencephalopathy with subcortical cysts: chronic white matter oedema due to a defect in brain ion and water homoeostasis. Lancet Neurol. 2012;11(11):973–85.

34. Rasmussen MK, Mestre H, and Nedergaard M. The glymphatic pathway in neurological disorders. Lancet Neurol. 2018;17(11):1016–24.

35. Owens GK. Regulation of differentiation of vascular smooth muscle cells. Physiol Rev. 1995;75(3):487–517.

36. Owens GK, Kumar MS, and Wamhoff BR. Molecular regulation of vascular smooth muscle cell differentiation in development and disease. Physiol Rev. 2004;84(3):767–801.

37. Chen ZL, Yao Y, Norris EH, Kruyer A, Jno-Charles O, Akhmerov A, et al. Ablation of astrocytic laminin impairs vascular smooth muscle cell function and leads to hemorrhagic stroke. J Cell Biol. 2013;202(2):381–95.

38. El-Khoury N, Braun A, Hu F, Pandey M, Nedergaard M, Lagamma EF, et al. Astrocyte end-feet in germinal matrix, cerebral cortex, and white matter in developing infants. Pediatr Res. 2006;59(5):673–9.

39. Takasato Y, Rapoport SI, and Smith QR. An in situ brain perfusion technique to study cerebrovascular transport in the rat. Am J Physiol. 1984;247(3 Pt 2):H484–93.

40. Anfray A, Drieu A, Hingot V, Hommet Y, Yetim M, Rubio M, et al. Circulating tPA contributes to neurovascular coupling by a mechanism involving the endothelial NMDA receptors. J Cereb Blood Flow Metab. 2019:271678X19883599.

41. Karlsson M, and Nordell B. Phantom and in vivo study of the Look-Locher T1 mapping method. Magn Reson Imaging. 1999;17(10):1481–8.

42. Gaberel T, Gakuba C, Goulay R, Martinez De Lizarrondo S, Hanouz JL, Emery E, et al. Impaired glymphatic perfusion after strokes revealed by contrast-enhanced MRI: a new target for fibrinolysis? Stroke. 2014;45(10):3092–6.

43. Schindelin J, Arganda-Carreras I, Frise E, Kaynig V, Longair M, Pietzsch T, et al. Fiji: an open-source platform for biological-image analysis. Nat Methods. 2012;9(7):676–82.

44. Renier N, Wu Z, Simon DJ, Yang J, Ariel P, and Tessier-Lavigne M. iDISCO: a simple, rapid method to immunolabel large tissue samples for volume imaging. Cell. 2014;159(4):896–910.

45. Pannasch U, Freche D, Dallerac G, Ghezali G, Escartin C, Ezan P, et al. Connexin 30 sets synaptic strength by controlling astroglial synapse invasion. Nat Neurosci. 2014;17(4):549–58.

46. Ghezali G, Calvo CF, Pillet LE, Llense F, Ezan P, Pannasch U, et al. Connexin 30 controls astroglial polarization during postnatal brain development. Development. 2018;145(4).

47. Bankhead P, Loughrey MB, Fernandez JA, Dombrowski Y, McArt DG, Dunne PD, et al. QuPath: Open source software for digital pathology image analysis. Sci Rep. 2017;7(1):16878.

