## Supplementary Figures and Tables for "Megalencephalic leukoencephalopathy with subcortical cysts is a developmental disorder of the gliovascular unit"

Fig. S1

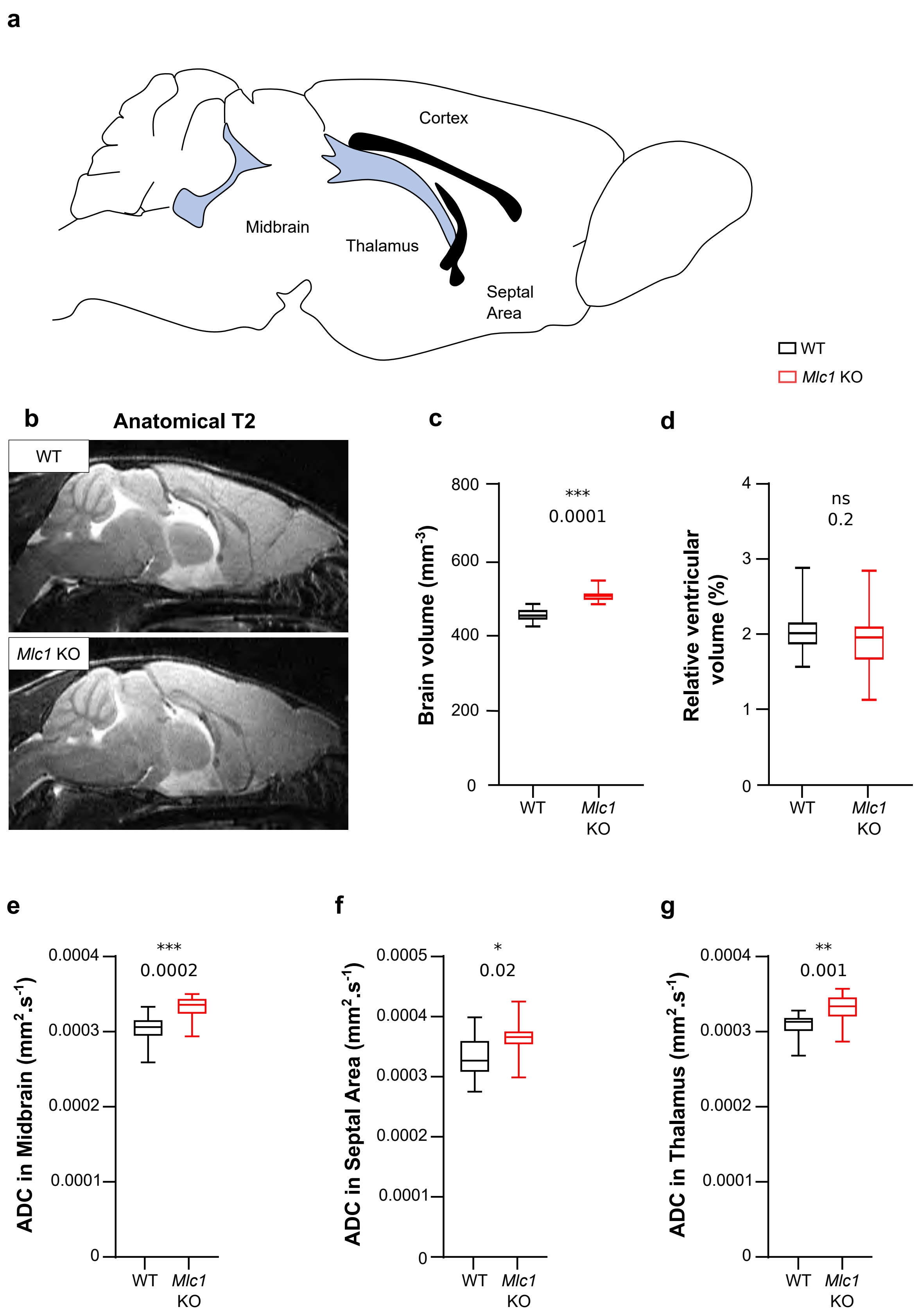
a
Cortex
Midbrain
Thalamus
Septal Area
WT
Mlc1 KO
b
Anatomical T2
WT
Mlc1 KO
c
***
0.0001
Brain volume (mm-3)
800
600
400
200
0
WT
Mlc1 KO
d
ns
0.2
Relative ventricular volume (%)
4
3
2
1
0
WT
Mlc1 KO
e
***
0.0002
ADC in Midbrain (mm2.s-1)
0.0004
0.0003
0.0002
0.0001
0
WT
Mlc1 KO
f
*
0.02
ADC in Septal Area (mm2.s-1)
0.0005
0.0004
0.0003
0.0002
0.0001
0
WT
Mlc1 KO
g
**
0.001
ADC in Thalamus (mm2.s-1)
0.0004
0.0003
0.0002
0.0001
0
WT
Mlc1 KO

Fig. S2

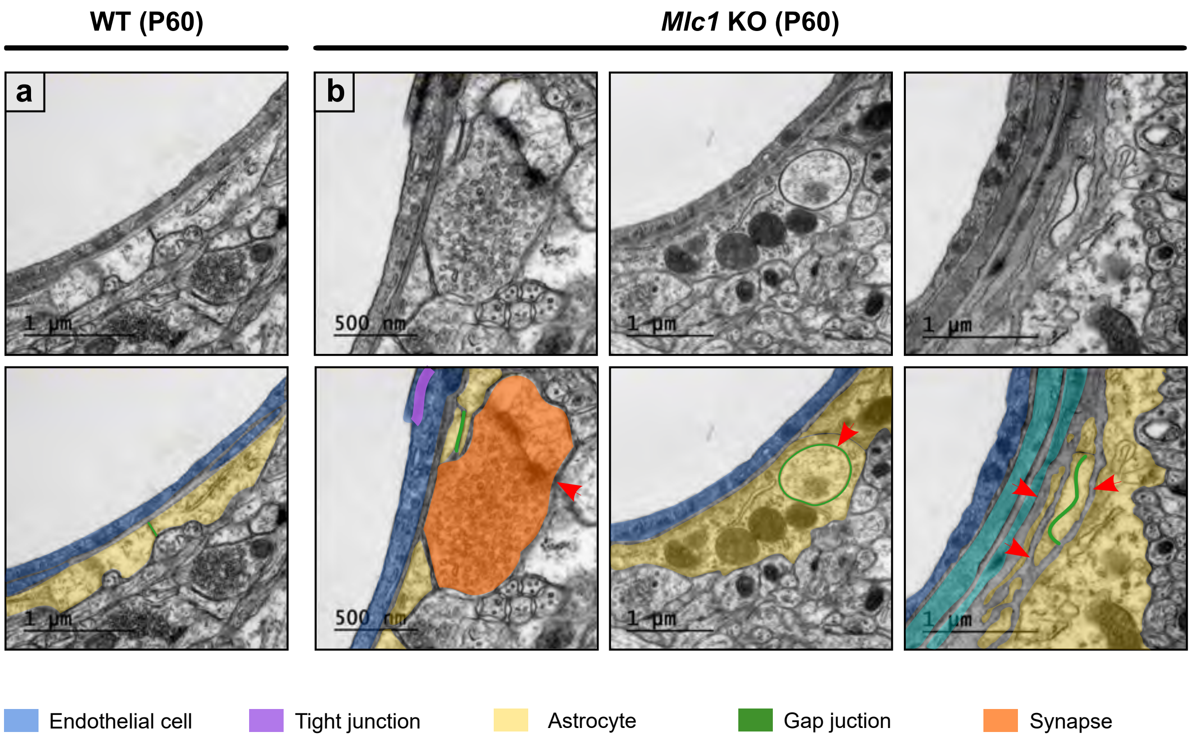

WT (P60)
*Mlc1* KO (P60)
a
b
1 µm
500 nm
1 µm
1 µm
1 µm
500 nm
1 µm
1 µm
Endothelial cell
Tight junction
Astrocyte
Gap juction
Synapse

Fig. S3

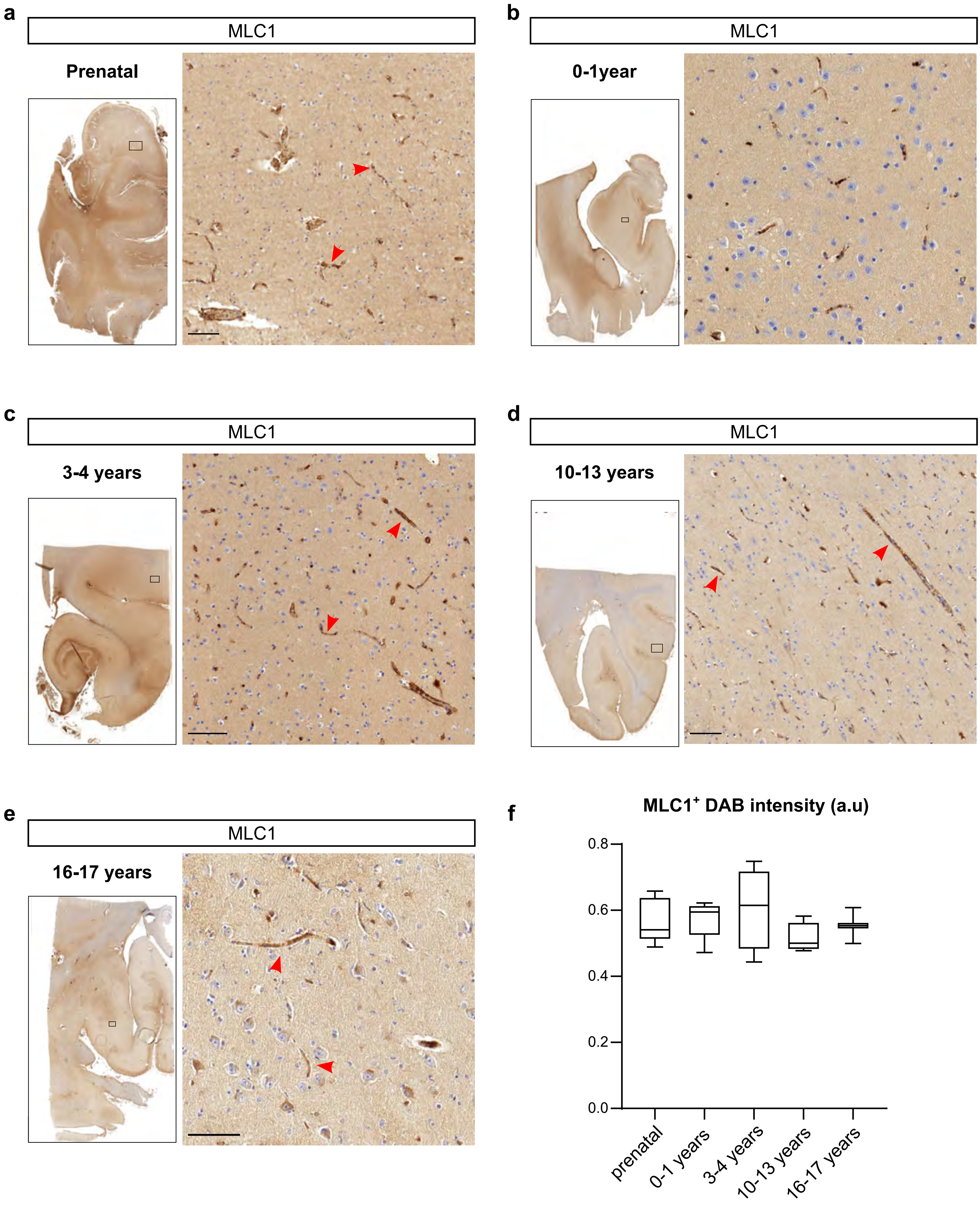
a
MLC1
Prenatal
b
MLC1
0-1year
c
MLC1
3-4 years
d
MLC1
10-13 years
e
MLC1
16-17 years
f
MLC1+ DAB intensity (a.u)
0.8
0.6
0.4
0.2
0.0
prenatal
0-1 years
3-4 years
10-13 years
16-17 years

### Table S1

endothelial cells (a.u.)

| Gene name | Stage | WT | | | | | | *Mlc1* KO | | | | | | Two-tailed Mann-Whitney test | |
|---|---|---|---|---|---|---|---|---|---|---|---|---|---|---|---|
| | | Min | 1st quartile | Median | 3rd quartile | Max | N | Min | 1st quartile | Median | 3rd quartile | Max | N | P-value | Significance |
| Abcb1 | P5 | 0.14 | 0.23 | 0.86 | 1.72 | 2.13 | 5 | 0.22 | 0.33 | 0.58 | 1.28 | 1.57 | 5 | >0.9999 | ns |
| | P15 | 10.38 | 11.16 | 7.39 | 15.42 | 16.03 | 4 | 6.64 | 6.73 | 7.09 | 10.30 | 11.34 | 4 | 0.0571 | ns |
| | P60 | 11.15 | 11.63 | 15.60 | 19.17 | 19.18 | 5 | 8.98 | 8.89 | 11.79 | 13.06 | 13.06 | 3 | 0.2500 | ns |
| Cldn5 | P5 | 2.38 | 3.01 | 5.95 | 7.96 | 8.28 | 4 | 6.25 | 7.10 | 10.14 | 13.49 | 14.45 | 4 | 0.1143 | ns |
| | P15 | 30.35 | 31.25 | 40.30 | 77.66 | 88.00 | 4 | 30.88 | 32.51 | 39.64 | 55.94 | 60.62 | 4 | >0.9999 | ns |
| | P60 | 8.61 | 9.35 | 13.83 | 19.16 | 20.18 | 4 | 11.58 | 11.65 | 16.98 | 27.16 | 28.85 | 4 | 0.3429 | ns |

VSMCs (a.u.)

| Gene name | Stage | WT | | | | | | *Mlc1* KO | | | | | | Two-tailed Mann-Whitney test | |
|---|---|---|---|---|---|---|---|---|---|---|---|---|---|---|---|
| | | Min | 1st quartile | Median | 3rd quartile | Max | N | Min | 1st quartile | Median | 3rd quartile | Max | N | P-value | Significance |
| Acta2 | P5 | 7.89 | 8.64 | 9.85 | 10.11 | 10.20 | 5 | 0.25 | 0.55 | 1.41 | 1.52 | 1.53 | 5 | 0.0079 | * |
| | P15 | 18.95 | 20.15 | 26.44 | 29.87 | 30.12 | 4 | 5.04 | 6.22 | 9.78 | 14.35 | 15.86 | 4 | 0.0286 | * |
| | P60 | 4.86 | 4.94 | 5.02 | 6.35 | 7.68 | 3 | 4.18 | 7.31 | 10.44 | 12.82 | 15.19 | 3 | 0.7000 | ns |
| Atp1b1 | P5 | 0.33 | 0.34 | 0.35 | 0.40 | 0.50 | 4 | 0.08 | 0.14 | 0.32 | 0.39 | 0.46 | 5 | 0.4127 | ns |
| | P15 | 0.19 | 0.29 | 0.35 | 0.42 | 0.58 | 4 | 0.40 | 0.51 | 0.61 | 0.68 | 0.70 | 4 | 0.1143 | ns |
| | P60 | 0.25 | 0.28 | 0.30 | 0.31 | 1.11 | 5 | 0.15 | 0.23 | 0.26 | 0.32 | 0.46 | 5 | 0.5476 | ns |

### Table S2

endothelial cells proteins (isolated GVU)

| Protein name | Stage | WT | | | | | | *Mlc1* KO | | | | | | Two-tailed Mann-Whitney test | |
|---|---|---|---|---|---|---|---|---|---|---|---|---|---|---|---|
| | | Min | 1st quartile | Median | 3rd quartile | Max | N | Min | 1st quartile | Median | 3rd quartile | Max | N | P-value | Significance |
| Pg-P | P5 | 0.40 | 0.55 | 0.58 | 0.70 | 0.91 | 5 | 0.30 | 0.42 | 0.47 | 0.49 | 0.50 | 4 | 0.1111 | ns |
| | P15 | 0.46 | 0.47 | 0.71 | 0.75 | 1.01 | 5 | 0.26 | 0.29 | 0.33 | 0.34 | 1.02 | 5 | 0.1508 | ns |
| | P60 | 0.35 | 0.57 | 0.87 | 0.88 | 1.26 | 5 | 0.68 | 0.70 | 0.77 | 0.97 | 1.09 | 5 | 0.8413 | ns |
| Claudin 5 | P5 | 0.26 | 0.49 | 0.59 | 0.73 | 0.85 | 5 | 0.49 | 0.54 | 0.59 | 0.63 | 0.79 | 5 | 0.8413 | ns |
| | P15 | 0.80 | 0.89 | 0.89 | 1.03 | 1.11 | 5 | 0.22 | 0.41 | 0.50 | 0.65 | 0.99 | 4 | 0.1111 | ns |
| | P60 | 1.16 | 1.27 | 1.37 | 1.41 | 1.51 | 5 | 1.05 | 1.22 | 1.32 | 1.38 | 1.54 | 5 | 0.8413 | ns |

VSMCs proteins (isolated GVU)

| Protein name | Stage | WT | | | | | | *Mlc1* KO | | | | | | Two-tailed Mann-Whitney test | |
|---|---|---|---|---|---|---|---|---|---|---|---|---|---|---|---|
| | | Min | 1st quartile | Median | 3rd quartile | Max | N | Min | 1st quartile | Median | 3rd quartile | Max | N | P-value | Significance |
| SMA | P5 | 0.12 | 0.14 | 0.21 | 0.38 | 0.80 | 5 | 0.16 | 0.20 | 0.46 | 0.65 | 1.90 | 5 | 0.4206 | ns |
| | P15 | 0.96 | 1.07 | 1.07 | 1.12 | 1.28 | 5 | 0.41 | 0.75 | 0.99 | 1.00 | 1.11 | 5 | 0.1508 | ns |
| | P60 | 1.15 | 1.16 | 1.20 | 1.38 | 1.78 | 4 | 0.69 | 0.97 | 1.04 | 1.05 | 1.15 | 5 | 0.0159 | * |

astrocytes protein (whole brain)(a.u.)

| Protein name | Stage | WT | | | | | | *Mlc1* KO | | | | | | Two-tailed Mann-Whitney test | |
|---|---|---|---|---|---|---|---|---|---|---|---|---|---|---|---|
| | | Min | 1st quartile | Median | 3rd quartile | Max | N | Min | 1st quartile | Median | 3rd quartile | Max | N | P-value | Significance |
| GlialCAM | P5 | 0.95 | 0.99 | 1.00 | 1.02 | 1.05 | 5 | 0.52 | 0.67 | 0.70 | 0.73 | 0.83 | 5 | 0.0079 | ** |
| | P15 | 0.35 | 0.58 | 0.96 | 1.10 | 2.01 | 5 | 0.17 | 0.29 | 0.72 | 0.84 | 1.62 | 5 | 0.4206 | ns |
| | P60 | 0.54 | 0.77 | 1.00 | 1.18 | 1.50 | 5 | 0.48 | 0.61 | 0.90 | 0.95 | 1.10 | 5 | 0.4206 | ns |
| Aqp 4 | P5 | 0.77 | 0.93 | 1.00 | 1.06 | 1.23 | 5 | 0.12 | 0.24 | 0.39 | 0.44 | 0.46 | 5 | 0.0079 | ** |
| | P15 | 0.47 | 0.84 | 0.89 | 1.31 | 1.49 | 5 | 0.72 | 0.80 | 0.88 | 0.92 | 0.98 | 5 | 0.6905 | ns |
| | P60 | 0.79 | 0.84 | 1.03 | 1.06 | 1.28 | 5 | 0.61 | 1.14 | 1.18 | 1.26 | 1.39 | 5 | 0.4206 | ns |
| Connexin 43 | P5 | 0.81 | 0.96 | 0.99 | 1.01 | 1.21 | 5 | 0.37 | 0.42 | 0.64 | 0.68 | 0.87 | 5 | 0.0159 | * |
| | P15 | 0.64 | 0.83 | 0.96 | 1.27 | 1.31 | 5 | 0.19 | 1.23 | 1.30 | 1.49 | 4.90 | 5 | 0.4206 | ns |
| | P60 | 0.66 | 0.92 | 0.96 | 1.20 | 1.26 | 5 | 0.63 | 1.48 | 1.58 | 1.98 | 2.10 | 5 | 0.1508 | ns |

astrocytes protein (MVs)(a.u.)

| Protein name | Stage | WT | | | | | | *Mlc1* KO | | | | | | Two-tailed Mann-Whitney test | |
|---|---|---|---|---|---|---|---|---|---|---|---|---|---|---|---|
| | | Min | 1st quartile | Median | 3rd quartile | Max | N | Min | 1st quartile | Median | 3rd quartile | Max | N | P-value | Significance |
| Aqp 4 | P60 | 0.48 | 0.52 | 1.08 | 1.44 | 1.71 | 5 | 0.26 | 0.27 | 0.28 | 0.46 | 0.47 | 5 | 0.0079 | ** |

neuronal protein (whole brain)(a.u.)

| Protein name | Stage | WT | | | | | | *Mlc1* KO | | | | | | Two-tailed Mann-Whitney test | |
|---|---|---|---|---|---|---|---|---|---|---|---|---|---|---|---|
| | | Min | 1st quartile | Median | 3rd quartile | Max | N | Min | 1st quartile | Median | 3rd quartile | Max | N | P-value | Significance |
| Neurofilament-M | P60 | 0.74 | 0.81 | 0.92 | 1.23 | 1.27 | 5 | 0.59 | 0.75 | 1.07 | 1.25 | 1.37 | 5 | 0.9999 | ns |

neuronal protein (MVs)(a.u.)

| Protein name | Stage | WT | | | | | | *Mlc1* KO | | | | | | Two-tailed Mann-Whitney test | |
|---|---|---|---|---|---|---|---|---|---|---|---|---|---|---|---|
| | | Min | 1st quartile | Median | 3rd quartile | Max | N | Min | 1st quartile | Median | 3rd quartile | Max | N | P-value | Significance |
| Neurofilament-M | P60 | 0.55 | 0.67 | 0.81 | 1.43 | 1.86 | 5 | 0.05 | 0.07 | 0.11 | 0.23 | 0.23 | 5 | 0.0079 | ** |

### Table S3

Volumetry

| | WT | | | | | | *Mlc1* KO | | | | | | Two-tailed Student's T test | |
|---|---|---|---|---|---|---|---|---|---|---|---|---|---|---|
| | Min | 1st quartile | Median | 3rd quartile | Max | N | Min | 1st quartile | Median | 3rd quartile | Max | N | P-value | Significance |
| Brain volume (mm3) | 423 | 443 | 452 | 469 | 483 | 17 | 483 | 494 | 505 | 511 | 545 | 18 | 0.0001 | *** |
| Relative ventricular volume (%) | 1.57 | 1.89 | 2.01 | 2.11 | 2.87 | 17 | 1.13 | 1.72 | 1.96 | 2.10 | 2.84 | 18 | 0.1975 | ns |

ADC (mm2.s-1)

| Zone | WT | | | | | | *Mlc1* KO | | | | | | Two-tailed Student's T test | |
|---|---|---|---|---|---|---|---|---|---|---|---|---|---|---|
| | Min | 1st quartile | Median | 3rd quartile | Max | N | Min | 1st quartile | Median | 3rd quartile | Max | N | P-value | Significance |
| Cerebellum | 0.00068 | 0.00078 | 0.00080 | 0.00083 | 0.00091 | 17 | 0.00077 | 0.00080 | 0.00083 | 0.00084 | 0.00086 | 17 | 0.3983 | ns |
| Cortex | 0.00078 | 0.00087 | 0.00090 | 0.00092 | 0.00098 | 17 | 0.00088 | 0.00097 | 0.00101 | 0.00103 | 0.00107 | 17 | 0.0001 | *** |
| Midbrain | 0.00078 | 0.00088 | 0.00092 | 0.00095 | 0.00100 | 17 | 0.00088 | 0.00097 | 0.00101 | 0.00103 | 0.00105 | 17 | 0.0002 | *** |
| Olfactory Bulb | 0.00078 | 0.00086 | 0.00090 | 0.00092 | 0.00097 | 17 | 0.00086 | 0.00090 | 0.00092 | 0.00094 | 0.00101 | 17 | 0.0557 | ns |
| Septal Area | 0.00083 | 0.00092 | 0.00098 | 0.00107 | 0.00120 | 17 | 0.00090 | 0.00106 | 0.00110 | 0.00113 | 0.00128 | 17 | 0.0210 | * |
| Thalamus | 0.00080 | 0.00091 | 0.00094 | 0.00096 | 0.00098 | 17 | 0.00086 | 0.00097 | 0.00100 | 0.00104 | 0.00107 | 17 | 0.0011 | ** |

Mean concentration of DOTA-Gd (µmol.mm-3)

| Zone | WT | | | | | | *Mlc1* KO | | | | | | Two-tailed Mann-Whitney test | |
|---|---|---|---|---|---|---|---|---|---|---|---|---|---|---|
| | Min | 1st quartile | Median | 3rd quartile | Max | N | Min | 1st quartile | Median | 3rd quartile | Max | N | P-value | Significance |
| Cerebellum | 0.0043 | 0.0267 | 0.0288 | 0.0484 | 0.0713 | 9 | 0.0254 | 0.0293 | 0.0467 | 0.0793 | 0.1031 | 9 | 0.1903 | ns |
| Cortex | 0.0000 | 0.0062 | 0.0078 | 0.0100 | 0.0167 | 8 | 0.0013 | 0.0045 | 0.0084 | 0.0095 | 0.0166 | 9 | 0.8148 | ns |
| Midbrain | 0.0047 | 0.0112 | 0.0259 | 0.0442 | 0.0660 | 9 | 0.0026 | 0.0086 | 0.0096 | 0.0146 | 0.0181 | 9 | 0.0315 | * |
| Olfactory Bulb | 0.0072 | 0.0114 | 0.0239 | 0.1220 | 0.1318 | 9 | 0.0046 | 0.0094 | 0.0118 | 0.0150 | 0.0246 | 9 | 0.1903 | ns |
| Septal Area | 0.0046 | 0.0146 | 0.0757 | 0.1286 | 0.1771 | 9 | 0.0071 | 0.0104 | 0.0118 | 0.0196 | 0.0321 | 9 | 0.1135 | ns |
| Thalamus | 0.0004 | 0.0051 | 0.0124 | 0.0273 | 0.0394 | 9 | 0.0014 | 0.0044 | 0.0052 | 0.0066 | 0.0170 | 9 | 0.1903 | ns |

Slope of DOTA-Gd (µmol.mm-3.min)

| Zone | WT | | | | | | *Mlc1* KO | | | | | | Two-tailed Mann-Whitney test | |
|---|---|---|---|---|---|---|---|---|---|---|---|---|---|---|
| | Min | 1st quartile | Median | 3rd quartile | Max | N | Min | 1st quartile | Median | 3rd quartile | Max | N | P-value | Significance |
| Cerebellum | 0.00009 | 0.00028 | 0.00054 | 0.00061 | 0.00139 | 9 | -0.00052 | -0.00019 | 0.00001 | 0.00011 | 0.00031 | 9 | 0.0012 | ** |
| Cortex | -0.00003 | 0.00001 | 0.00014 | 0.00022 | 0.00029 | 8 | -0.00014 | -0.00007 | -0.00007 | 0.00007 | 0.00009 | 9 | 0.0152 | * |
| Midbrain | 0.00009 | 0.00022 | 0.00053 | 0.00100 | 0.00124 | 9 | -0.00008 | 0.00005 | 0.00013 | 0.00026 | 0.00044 | 9 | 0.0188 | * |
| Olfactory Bulb | -0.00071 | -0.00003 | 0.00013 | 0.00038 | 0.00100 | 9 | -0.00005 | -0.00003 | 0.00009 | 0.00018 | 0.00042 | 9 | 0.9314 | ns |
| Septal Area | -0.00040 | -0.00002 | 0.00020 | 0.00021 | 0.00128 | 9 | -0.00022 | 0.00002 | 0.00006 | 0.00016 | 0.00063 | 9 | 0.5457 | ns |
| Thalamus | -0.00008 | 0.00004 | 0.00047 | 0.00081 | 0.00107 | 9 | -0.00010 | 0.00004 | 0.00008 | 0.00010 | 0.00023 | 9 | 0.0770 | ns |

### Table S4

BBB permeability (µL/g)

| | WT | | | | | | *Mlc1* KO | | | | | | Two-tailed Mann-Whitney test | |
|---|---|---|---|---|---|---|---|---|---|---|---|---|---|---|
| | Min | 1st quartile | Median | 3rd quartile | Max | N | Min | 1st quartile | Median | 3rd quartile | Max | N | P-value | Significance |
| Albumin - | 10.00 | 14.18 | 16.75 | 18.73 | 24.50 | 8 | 9.20 | 14.35 | 15.70 | 18.60 | 26.50 | 9 | 0.9824 | ns |
| Albumin + | 12.00 | 13.40 | 14.80 | 18.00 | 22.40 | 11 | 10.00 | 12.15 | 13.95 | 19.38 | 30.00 | 12 | 0.4399 | ns |

### Table S5

| Pecam-1 | | WT | | | | | | *Mlc1* KO | | | | | | One-tailed Mann-Whitney test | |
|---|---|---|---|---|---|---|---|---|---|---|---|---|---|---|---|
| | | Min | 1st quartile | Median | 3rd quartile | Max | N | Min | 1st quartile | Median | 3rd quartile | Max | N | P-value | Significance |
| Parenchymal | length ratio (µm-2) | 1.72E-05 | 2.00E-05 | 2.28E-05 | 2.34E-05 | 2.41E-05 | 3 | 1.39E-05 | 1.66E-05 | 1.93E-05 | 1.92E-04 | 3.65E-04 | 3 | 0.3500 | ns |
| | branching ratio (µm-3) | 2.79E-05 | 3.23E-05 | 3.66E-05 | 3.90E-05 | 4.15E-05 | 3 | 2.29E-05 | 2.74E-05 | 3.20E-05 | 4.55E-05 | 5.89E-05 | 3 | 0.5000 | ns |
| | straightness (µm-3) | 9.40E-01 | 9.45E-01 | 9.50E-01 | 9.50E-01 | 9.50E-01 | 3 | 9.40E-01 | 9.40E-01 | 9.40E-01 | 9.50E-01 | 9.60E-01 | 3 | 0.5000 | ns |
| Cortical surface | length ratio (µm-1) | 9.17E-05 | 1.07E-04 | 1.22E-04 | 1.90E-04 | 2.58E-04 | 3 | 5.52E-05 | 8.76E-05 | 1.20E-04 | 1.23E-04 | 1.25E-04 | 3 | 0.3500 | ns |
| | branching ratio (µm-2) | 1.33E-06 | 1.95E-06 | 2.58E-06 | 4.02E-06 | 5.45E-06 | 3 | 5.96E-07 | 1.51E-06 | 2.43E-06 | 2.50E-06 | 2.57E-06 | 3 | 0.2000 | ns |
| | straightness (µm-2) | 9.20E-01 | 9.25E-01 | 9.30E-01 | 9.35E-01 | 9.40E-01 | 3 | 9.30E-01 | 9.35E-01 | 9.40E-01 | 9.45E-01 | 9.50E-01 | 3 | 0.2500 | ns |

| SMA | | WT | | | | | | *Mlc1* KO | | | | | | One-tailed Mann-Whitney test | |
|---|---|---|---|---|---|---|---|---|---|---|---|---|---|---|---|
| | | Min | 1st quartile | Median | 3rd quartile | Max | N | Min | 1st quartile | Median | 3rd quartile | Max | N | P-value | Significance |
| parenchymal | length ratio (µm-2) | 1.63E-05 | 2.00E-05 | 2.37E-05 | 2.43E-05 | 2.49E-05 | 3 | 1.86E-05 | 2.52E-05 | 3.18E-05 | 4.93E-05 | 6.69E-05 | 3 | 0.2000 | ns |
| | branching ratio (µm-3) | 1.85E-07 | 2.65E-07 | 3.45E-07 | 3.45E-07 | 3.45E-07 | 3 | 2.75E-07 | 3.15E-07 | 3.55E-07 | 6.06E-07 | 8.57E-07 | 3 | 0.2000 | ns |
| | straightness (µm-3) | 9.12E-01 | 9.23E-01 | 9.33E-01 | 9.34E-01 | 9.34E-01 | 3 | 8.95E-01 | 9.13E-01 | 9.32E-01 | 9.49E-01 | 9.65E-01 | 3 | 0.5000 | ns |
| cortical surface | length ratio (µm-1) | 1.63E-03 | 1.83E-03 | 1.91E-03 | 1.97E-03 | 2.14E-03 | 4 | 7.66E-04 | 1.13E-03 | 1.47E-03 | 1.76E-03 | 1.95E-03 | 4 | 0.1714 | ns |
| | branching ratio (µm-2) | 7.48E-06 | 8.70E-06 | 1.56E-05 | 2.33E-05 | 2.73E-05 | 4 | 5.21E-06 | 5.74E-06 | 8.71E-06 | 1.44E-05 | 2.30E-05 | 4 | 0.2429 | ns |
| | straightness (µm-2) | 8.99E-01 | 9.11E-01 | 9.16E-01 | 9.19E-01 | 9.27E-01 | 4 | 9.07E-01 | 9.11E-01 | 9.16E-01 | 9.23E-01 | 9.30E-01 | 4 | 0.4429 | ns |
| | anastomosis (µm-2) | 2.10E-07 | 2.28E-07 | 2.80E-07 | 3.70E-07 | 5.00E-07 | 4 | 1.14E-07 | 1.77E-07 | 2.80E-07 | 3.75E-07 | 4.13E-07 | 4 | 0.3429 | ns |

### Table S6

amplitude (%.min-1)

| | WT | | | | | | *Mlc1* KO | | | | | | Two-tailed Mann-Whitney test | |
|---|---|---|---|---|---|---|---|---|---|---|---|---|---|---|
| | Min | 1st quartile | Median | 3rd quartile | Max | N | Min | 1st quartile | Median | 3rd quartile | Max | N | P-value | Significance |
| P5 | -0.38 | 3.13 | 4.55 | 8.14 | 16.78 | 13.00 | -0.84 | 1.08 | 2.35 | 3.33 | 5.97 | 13 | 0.0501 | ns |
| P10 | 3.73 | 8.59 | 13.65 | 16.74 | 29.07 | 9.00 | 1.45 | 3.70 | 5.26 | 9.51 | 11.87 | 12 | 0.0073 | ** |
| P15 | 3.55 | 11.96 | 14.58 | 10.75 | 48.06 | 8.00 | -0.63 | 9.87 | 12.26 | 13.37 | 24.42 | 11 | 0.0091 | ** |

slope (%)

| | WT | | | | | | *Mlc1* KO | | | | | | Two-tailed Mann-Whitney test | |
|---|---|---|---|---|---|---|---|---|---|---|---|---|---|---|
| | Min | 1st quartile | Median | 3rd quartile | Max | N | Min | 1st quartile | Median | 3rd quartile | Max | N | P-value | Significance |
| P5 | -3.95 | -1.30 | -0.72 | -0.31 | 0.27 | 13.00 | -1.75 | -0.72 | -0.50 | -0.20 | 0.74 | 13 | 0.2035 | ns |
| P10 | -3.04 | -2.65 | -1.05 | -1.01 | -0.28 | 9.00 | -2.03 | -1.40 | -1.15 | -0.61 | -0.35 | 12 | 0.0339 | * |
| P15 | -5.32 | -3.64 | -3.39 | -2.79 | -0.42 | 8.00 | -4.13 | -3.00 | -2.02 | -0.53 | 0.07 | 11 | 0.0067 | ** |

### Table S7

Microbubbles Ultrafast Ultrasound - arterial diameter (µm)

| WT | | | | | | *Mlc1* KO | | | | | | Two-tailed Mann-Whitney test | |
|---|---|---|---|---|---|---|---|---|---|---|---|---|---|
| Min | 1st quartile | Median | 3rd quartile | Max | N | Min | 1st quartile | Median | 3rd quartile | Max | N | P-value | Significance |
| 36.18 | 37.34 | 38.30 | 38.83 | 44.18 | 6 | 29.67 | 31.31 | 32.07 | 33.50 | 33.89 | 6 | 0.0022 | ** |

Fonctionnal  Ultrafast Ultrasound - % of Power Doppler increase

| WT | | | | | | *Mlc1* KO | | | | | | Two-tailed Mann-Whitney test | |
|---|---|---|---|---|---|---|---|---|---|---|---|---|---|
| Min | 1st quartile | Median | 3rd quartile | Max | N | Min | 1st quartile | Median | 3rd quartile | Max | N | P-value | Significance |
| 5.22 | 5.96 | 7.46 | 8.10 | 13.84 | 11 | 2.28 | 2.61 | 3.86 | 5.33 | 10.36 | 12 | 0.0028 | ** |

### Table S8

Sholl (a.u.)

| Stage | Distance to the soma (mm) | WT | | | | | | *Mlc1* KO | | | | | | Bonferroni's multiple comparaison | | 2-way-Anova | |
|---|---|---|---|---|---|---|---|---|---|---|---|---|---|---|---|---|---|
| | | Min | 1st quartile | Median | 3rd quartile | Max | N | Min | 1st quartile | Median | 3rd quartile | Max | N | P-value | Signifiance | P-value | Signifiance |
| P10 | 10.00 | 4 | 7 | 9 | 11 | 14 | 49 | 4 | 10 | 12 | 13 | 24 | 53 | 0.0001 | *** | 0.0001 | *** |
| | 15.00 | 5 | 9 | 9 | 12 | 17 | | 7 | 11 | 13 | 16 | 26 | | 0.0001 | *** | | |
| | 20.00 | 2 | 6 | 8 | 11 | 22 | | 5 | 8 | 10 | 14 | 19 | | 0.0001 | *** | | |
| | 25.00 | 0 | 3 | 6 | 7 | 18 | | 0 | 3 | 5 | 9 | 16 | | 0.9999 | ns | | |
| | 30.00 | 0 | 1 | 3 | 4 | 9 | | 0 | 1 | 2 | 4 | 7 | | 0.9999 | ns | | |
| | 35.00 | 0 | 0 | 0 | 2 | 6 | | 0 | 0 | 0 | 1 | 5 | | 0.9999 | ns | | |
| P15 | 10.00 | 3 | 6 | 8 | 10 | 20 | 50 | 7 | 11 | 13 | 16 | 21 | 51 | 0.0001 | *** | 0.0001 | *** |
| | 15.00 | 3 | 7 | 10 | 13 | 24 | | 4 | 11 | 15 | 19 | 23 | | 0.0001 | *** | | |
| | 20.00 | 2 | 5 | 9 | 11 | 17 | | 6 | 10 | 11 | 16 | 21 | | 0.0001 | *** | | |
| | 25.00 | 0 | 3 | 5 | 8 | 14 | | 2 | 6 | 7 | 10 | 18 | | 0.0005 | *** | | |
| | 30.00 | 0 | 1 | 2 | 4 | 12 | | 0 | 2 | 4 | 7 | 10 | | 0.3544 | ns | | |
| | 35.00 | 0 | 0 | 0 | 2 | 6 | | 0 | 0 | 1 | 2 | 7 | | 0.9999 | ns | | |
| P60 | 10.00 | 7 | 11 | 14 | 17 | 27 | 48 | 6 | 10 | 12 | 14 | 21 | 44 | 0.6570 | ns | 0.0001 | *** |
| | 15.00 | 6 | 13 | 18 | 21 | 27 | | 7 | 11 | 14 | 16 | 22 | | 0.0003 | *** | | |
| | 20.00 | 7 | 12 | 16 | 19 | 26 | | 6 | 10 | 13 | 17 | 30 | | 0.3030 | * | | |
| | 25.00 | 1 | 6 | 10 | 13 | 23 | | 2 | 6 | 7 | 9 | 13 | | 0.0011 | ** | | |
| | 30.00 | 0 | 2 | 5 | 8 | 16 | | 0 | 2 | 3 | 5 | 9 | | 0.3927 | ns | | |
| | 35.00 | 0 | 0 | 1 | 3 | 8 | | 0 | 0 | 1 | 2 | 6 | | 0.9999 | ns | | |

Orientation to the stratum radiatum (a.u.)

| Stage | WT | | | | | | One sample wilcoxon test | | *Mlc1* KO | | | | | | One sample wilcoxon test | | Two-tailed Mann-Whitney test | |
|---|---|---|---|---|---|---|---|---|---|---|---|---|---|---|---|---|---|---|
| | Min | 1st quartile | Median | 3rd quartile | Max | N | P-value | Signifiance | Min | 1st quartile | Median | 3rd quartile | Max | N | P-value | Signifiance | P-value | Significance |
| P60 | 0.63 | 1.05 | 1.20 | 1.62 | 3.18 | 47.00 | 0.0001 | *** | 0.47 | 1.07 | 1.19 | 1.56 | 2.45 | 52.00 | 0.0001 | *** | 0.8904 | ns |

Orientation to vessels (a.u.)

| Stage | WT | | | | | | One sample wilcoxon test | | *Mlc1* KO | | | | | | One sample wilcoxon test | | Two-tailed Mann-Whitney test | |
|---|---|---|---|---|---|---|---|---|---|---|---|---|---|---|---|---|---|---|
| | Min | 1st quartile | Median | 3rd quartile | Max | N | P-value | Signifiance | Min | 1st quartile | Median | 3rd quartile | Max | N | P-value | Signifiance | P-value | Significance |
| P10 | 0.50 | 0.85 | 1.03 | 1.14 | 1.69 | 26.00 | 0.7710 | ns | 0.33 | 1.00 | 1.19 | 1.28 | 1.73 | 35.00 | 0.0072 | ** | 0.0401 | * |
| P15 | 0.67 | 0.90 | 1.10 | 1.14 | 1.50 | 46.00 | 0.1157 | ns | 0.62 | 1.00 | 1.15 | 1.36 | 1.78 | 48.00 | 0.0004 | *** | 0.0482 | * |
| P60 | 0.56 | 0.88 | 1.03 | 1.11 | 1.59 | 41.00 | 0.8225 | ns | 0.91 | 1.13 | 1.22 | 1.42 | 2.29 | 41.00 | 0.0001 | *** | 0.0001 | *** |

### Table S9

% of capillaries and venules with contacting neuronal processes :

| Age | zone | WT | | | | | | *Mlc1* KO | | | | | | Two-tailed Student's T test | |
|---|---|---|---|---|---|---|---|---|---|---|---|---|---|---|---|
| | | Min | 1st quartile | Median | 3rd quartile | Max | N (mice) | Min | 1st quartile | Median | 3rd quartile | Max | N (mice) | P-value | Significance |
| P60 | Cortex | 5.95 | 7.37 | 8.15 | 9.15 | 11.25 | 4 | 26.36 | 43.66 | 51.46 | 54.99 | 59.48 | 4 | 0.0018 | *** |
| | Hippocampus | 4.62 | 4.78 | 5.54 | 6.25 | 6.25 | 4 | 19.32 | 21.02 | 25.80 | 30.68 | 32.73 | 4 | 0.0008 | *** |

Swelling :

| Age | zone | | WT | | | | | | *Mlc1* KO | | | | | | Two-tailed Mann-Whitney test | |
|---|---|---|---|---|---|---|---|---|---|---|---|---|---|---|---|---|
| | | | Min | 1st quartile | Median | 3rd quartile | Max | N (mice) | Min | 1st quartile | Median | 3rd quartile | Max | N (mice) | P-value | Significance |
| P60 | Cortex | swelling 0 | 56.30 | 74.00 | 81.15 | 83.75 | 87.80 | 4 | 45.80 | 53.00 | 68.40 | 81.73 | 82.70 | 4 | 0.2429 | ns |
| | | swelling 1 | 12.20 | 15.43 | 18.30 | 25.70 | 42.50 | 4 | 17.30 | 18.28 | 31.20 | 45.93 | 52.30 | 4 | 0.1714 | ns |
| | | swelling 2 | 0.00 | 0.00 | 0.55 | 1.13 | 1.20 | 4 | 0.00 | 0.00 | 0.45 | 1.18 | 2.00 | 4 | 0.5000 | ns |
| | Hippocampus | swelling 0 | 65.60 | 67.85 | 71.70 | 78.35 | 89.00 | 4 | 51.90 | 55.73 | 64.60 | 74.30 | 80.60 | 4 | 0.2429 | ns |
| | | swelling 1 | 11.00 | 21.65 | 28.30 | 32.00 | 33.80 | 4 | 18.50 | 25.48 | 32.65 | 40.15 | 48.10 | 4 | 0.2429 | ns |
| | | swelling 2 | 0.00 | 0.00 | 0.00 | 0.15 | 0.60 | 4 | 0.00 | 0.00 | 0.40 | 1.98 | 5.50 | 4 | 0.2143 | ns |

Absence of glial coverage (%) :

| Age | zone | WT | | | | | | *Mlc1* KO | | | | | | Two-tailed Mann-Whitney test | |
|---|---|---|---|---|---|---|---|---|---|---|---|---|---|---|---|
| | | Min | 1st quartile | Median | 3rd quartile | Max | N (mice) | Min | 1st quartile | Median | 3rd quartile | Max | N (mice) | P-value | Significance |
| P5 | Cortex | 41.04 | 41.38 | 41.73 | 58.54 | 50.13 | 3 | 37.67 | 39.91 | 46.59 | 54.53 | 64.55 | 4 | 0.8959 | ns |
| P10 | | 94.26 | 94.83 | 95.40 | 97.09 | 96.25 | 3 | 84.53 | 85.85 | 87.18 | 88.49 | 89.80 | 3 | 0.0001 | *** |
| P15 | | 98.13 | 98.73 | 99.06 | 99.55 | 99.27 | 4 | 93.18 | 93.40 | 94.01 | 94.94 | 96.11 | 4 | 0.0001 | *** |
| P60 | | 99.90 | 99.91 | 99.91 | 99.94 | 99.93 | 3 | 95.91 | 96.19 | 96.47 | 96.63 | 96.79 | 3 | 0.0001 | *** |

| Two-tailed Mann-Whitney test | | |
|---|---|---|
| Groups | P-value | Signifiance |
| WT P5/WT 10 | 0.0001 | *** |
| WT P10/WT P15 | 0.0001 | *** |
| WT P15/WT P60 | 0.0001 | *** |

### Table S10

MLC1 + DAB intentity (a.u)

| | Min | 1st quartile | Median | 3rd quartile | Max | N | Two-tailed Mann-Whitney test | | |
|---|---|---|---|---|---|---|---|---|---|
| | | | | | | | Groups | P-value | Signifiance |
| Prenatal | 0.4890 | 0.5311 | 0.5412 | 0.6236 | 0.6583 | 5 | prenatal/0-1 year | 0.9999 | ns |
| 0-1 year | 0.4721 | 0.5721 | 0.5951 | 0.6104 | 0.6224 | 5 | prenatal/3-5 years | 0.7302 | ns |
| 3-5 years | 0.4436 | 0.4795 | 0.5373 | 0.6273 | 0.7482 | 4 | prenatal/10-13 years | 0.1904 | ns |
| 10-13 years | 0.4778 | 0.4847 | 0.5163 | 0.5825 | 0.6083 | 4 | 0-1 year/3-5 years | 0.7302 | ns |
| 16-17 years | 0.4994 | / | / | / | 0.6083 | 2 | 0-1 year/ 10-13 years | 0.2858 | ns |
| | | | | | | | 3-5 years/10-13 years | 0.3428 | ns |

### Table S11

qPCR primers

| Gene | Forward/Reverse | Sequence |
|---|---|---|
| Acta2 | F | GTCCCAGACATCAGGGAGTAA |
| Acta2 | R | TCGGATACTTCAGCGTCAGGA |
| Atp1b1 | F | GCTGCTAACCATCAGTGAACT |
| Atp1b1 | R | GGGGTCATTAGGACGGAAGGA |
| Mdr1a (Abcb1) | F | GATAGGCTGGTTTGATGTGC |
| Mdr1a (Abcb1) | R | TCACAAGGGTTAGCTTCCAG |
| Cldn5 | F | TAAGGCACGGGTAGCACTCA |
| Cldn5 | R | GGACAACGATGTTGGCGAAC |
| Gapdh | F | AGGTCGGTGTGAACGGATTTG |
| Gapdh | R | TGTAGACCATGTAGTTGAGGTCA |

Antibodies

| | Protein / molecule | Species | Reference | Supplier | Western blot dilution | Immunohistology dilution |
|---|---|---|---|---|---|---|
| Primary antibodies | Claudin 5 | rabbit | 34-1600 | THERMO FISHER | 1:500 | / |
| | H3 | mouse | 14269S | OZYME | 1:2000 | / |
| | P-gP | mouse | ALX-801-002-C100 | ENZO | 1:200 | / |
| | SMA_cy3 | mouse | C6198 | SIGMA | 1:1000 | 1:250 |
| | Pecam-1 | goat | AF3628 | R&D SYSTEMS | / | 1:300 |
| | Connexin 43 | mouse | 610062 | BD transduction lab | 1:500 | 1:200 |
| | GlialCAM | rabbit | / | Provided by Raul Estevez, Universitat de Barcelona, Spain. | 1:500 | 1:500 |
| | Aquaporin 4 | rabbit | A5971 | SIGMA | 1:500 | 1:500 |
| | Neurofilament M | mouse | / | Provided by Dr Beat M. Riederer, University of Lausanne, Switzerland. | 1:40 | / |
| | GFAP | rabbit | G9269 | SIGMA | / | 1:500 |
| | MLC1human | rabbit | / | Provided by Raul Estevez, Universitat de Barcelona, Spain. | / | 1:200 |
| Secondary antibodies | anti-rabbit_alexa 555 | goat | A21429 | INVITROGEN | / | 1:2000 |
| | anti-rabbit_alexa 488 | goat | A11034 | INVITROGEN | / | 1:2000 |
| | anti-mouse_alexa 555 | goat | A21424 | INVITROGEN | / | 1:2000 |
| | anti-goat_alexa 647 | donkey | A-21447 | LIFE | / | 1:2000 |
| | anti-rabbit_HRP | goat | CSA2115 | COHESION | 1:2500 | / |
| | anti-mouse_HRP | goat | CSA2108 | COHESION | 1:2500 | / |
| Other molecules | IB4 | Alexa-conjugated Isolectin (*griffonia simplicifolia* ) | I32450 | INVITROGEN | 1:100 | / |
